# Sex differences in epigenetic mechanisms of chronic pain-induced depression

**DOI:** 10.64898/2026.07.04.736250

**Authors:** Mithil Gaikwad, Vandana Shree Vedartham Srinivasan, Beyza Ayazgok, Mathieu Bruggeman, Heba Elseedy, Hanus Slavik, Alba Caparros-Roissard, Yahia Hadj-Arab, Khaled Abdallah, Noémie Willem, Stéphanie Le Gras, Benoit Labonté, Ipek Yalcin, Pierre-Eric Lutz

## Abstract

Chronic pain is a major risk factor for depression, yet the molecular mechanisms underlying this comorbidity remain poorly understood, particularly in women. To address this gap, we systematically investigated sex differences in the epigenomic adaptations associated with chronic pain-induced depressive-like behaviors. Neuropathic pain was induced in the mouse using the sciatic nerve cuff model, and molecular analyses were performed in the anterior cingulate cortex (ACC), a key brain region implicated in both pain and affective processing. We profiled genome-wide DNA methylation, three histone modifications (H3K27ac, H3K4me1, and H3K27me3), and gene expression using EM-seq, Cut&Tag sequencing, and RNA-seq, respectively. Differential analyses were conducted for each molecular layer and integrated through gene co-expression network analysis. We found that chronic pain induced extensive remodeling of DNA methylation and histone modification landscapes in both sexes. Strikingly, these changes occurred at largely distinct genomic loci in males and females, revealing pronounced sex-specific epigenetic responses. Despite this divergence, the affected regions displayed similar regulatory organization, including enrichment at shared genic features, transcription factor binding sites, and chromatin profiles. Importantly, these adaptations converged on partly overlapping genes, biological pathways, and co-expression modules across sexes. The most affected gene modules were predominantly associated with synapse-related processes, consistent with previous knowledge, and were closely connected to modules enriched for epigenetic regulatory functions. Together, these findings indicate that chronic pain engages sex-specific epigenetic mechanisms that ultimately converge on common functional outcomes. Such convergence highlights the potential value of targeting sex-specific epigenetic substrates in future therapeutic strategies.

## Introduction

Chronic neuropathic pain is a debilitating condition that impacts approximately 7 to 10% of the general population over the lifetime [1]. Up to 40-60% [2] of patients with chronic neuropathic pain also develop an anxiety or depressive disorder [3, 4], resulting in frequent comorbidities that associate with a poorer prognosis [5]. Therefore, understanding the molecular mechanisms underlying these long-term consequences is important to improve pain management.

Numerous data indicate that there are notable sex differences in the prevalence, manifestation, and progression of chronic pain [6–8]. Epidemiological studies show that women are more frequently affected by certain types of pain, including neuropathic pain, and exhibit a higher susceptibility to comorbid anxiety or depression, compared to men [9, 10]. Sex differences have also been reliably observed at clinical level, using both quantitative sensory testing and evaluation of psychiatric symptomatology [11–14]. Surprisingly, the molecular mechanisms that may account for such differences have only recently started to be characterized in preclinical models. Among other findings, available studies indicate that male and female rodents show distinct changes in gene expression [15–18], receptor signaling, or activity of specific neuronal circuits [19] during chronic pain. However, how these sex-specific adaptations are initiated at the genomic level remains poorly defined.

Epigenetic mechanisms represent promising candidates to help explain the genomic reprogramming recruited during neuropsychiatric conditions. These mechanisms, which include DNA methylation and histone post-translational modifications [20], can be modulated by life experiences [21] and have been proposed to mediate long-lasting changes in neural function and behavior [22, 23]. More specifically, they have been implicated in the modulation of both nociception [24] and mood [25, 26], and may therefore contribute to the pain-depression comorbidity. Studies conducted on DNA methylation in rodent models of pain focused on a limited set of genes (*Kcna2, Oprm1, Dnmt*) [27–34], with only one report that used microarrays to more broadly analyze promoters [35]. The majority of the DNA methylome therefore remains unexplored, in particular outside promoters, where most plasticity likely occurs [36]. Similarly, studies on histone modifications focused on candidate genes, and only 2 reports investigated a single histone mark in the whole genome (H3K9ac [37], H3K27me3 [38]). Overall, how chronic pain may differentially affect multiple omic layers, and dysregulate their cross-talk, remains unknown.

To address these gaps, we conducted a genome-wide characterization of the multiomic plasticity that is recruited when pain leads to mood dysfunction, and systematically assessed how this plasticity differs as a function of sex [39]. Building on our mouse model of chronic neuropathic pain-induced depressive-like behaviors (CPID) [40], we used high-throughput approaches to analyze the anterior cingulate cortex (ACC), a major brain structure for this comorbidity [41, 42]. DNA methylation and 3 histone modifications (H3K27ac, H3K27me3, and H3K4me1) were assessed using Enzymatic Methylation sequencing (EM-seq) and Cleavage Under Targets and Tagmentation sequencing (CUT&Tag), respectively. We also defined the transcriptional impact of the observed epigenetic changes, using RNA-Sequencing. Finally, we leveraged gene network theory to define the organization of gene co-expression, and prioritize groups of genes, defined as modules, most significantly affected by pain-induced molecular changes.

Briefly, results indicated that CPID was associated with DNA methylation and histone changes that occurred at highly distinct genomic loci across females and males, unraveling sex-specific forms of epigenomic plasticity. These multiomic changes nevertheless affected strikingly similar gene modules and biological functions, revealing a potent functional convergence. Dysregulated pathways notably included synaptic plasticity, consistent with previous knowledge [28, 43], as well as the epigenetic machinery itself, highlighting its contribution to chronic pain. These adaptations were primarily enriched for genes expressed by excitatory and inhibitory neurons, indicating that these cells may be particularly sensitive to pain-induced epigenetic changes. Overall, the present work identifies epigenetic targets that, in the long term, may be modulated in a sex-specific manner to more effectively alleviate the negative emotional consequences of pain.

## Materials and Methods

See also supplementary material.

### Animals

We used male and female C57BL/6J mice (Charles River, l’Arbresle, France) aged 8 weeks upon arrival. Mice were group housed (5 per cage) with enrichment and maintained on a 12-hr light-dark cycle (lights on: 8pm and off 8 am) at 24±1°C, humidity around 50%, with *ad libitum* access to food and water.

### Neuropathic pain model

Following 2 weeks of habituation, baseline mechanical nociceptive thresholds were evaluated prior to surgery. Then, neuropathic pain was induced as previously described [40, 44]. A 2 mm polyethylene cuff (Harvard Apparatus, Les Ulis, France) was placed around the main branch of the right sciatic nerve, under general anesthesia (ketamine 100 mg/kg, xylazine 10 mg/kg; Centravet, Taden, France). Mice from the sham control group underwent an identical surgical procedure but without any cuff implantation.

## Results

### Chronic Pain-Induced Depression (CPID) triggers widespread DNA methylation changes in male and female mice

Over the years, we have extensively used a mouse model of chronic neuropathic pain induced by implantation of a polyethylene “cuff” around the main branch of the sciatic nerve. While this model was initially established in males [40, 45, 46], we more recently extended it to females by demonstrating that they similarly develop depressive-like behaviors following neuropathic pain induction (Vedartham Srinivasan et al, In preparation). Based on these results, here we processed cohorts of male and female mice in this model (Fig.1a; N=6/sex/group, 24 total). As expected, a long-lasting mechanical hypersensitivity was observed in both sexes in CPID mice compared to Sham controls (Males: (*F*(14.3, 194.0)=5.8, p< 0.0001; post-hoc 1-7weeks, p<0.05); Females: (*F*(18.0, 240.3)=3.5, p<0.0001; post-hoc 1-8 weeks, p<0.05) (Fig.1b). This hypersensitivity was accompanied, 8 weeks after neuropathy induction, by depressive-like behaviors in the Splash test (Males: Mann-Whitney *U*=26, *p*=0.0066; Females: Welch’s *t*(18.16)= 8.057, *p*<0.0001) in both sexes (Fig.1c). At this timepoint, ACC tissue was dissected from these cohorts, and used for an array of epigenomic and transcriptomic investigations.

**Figure 1.**
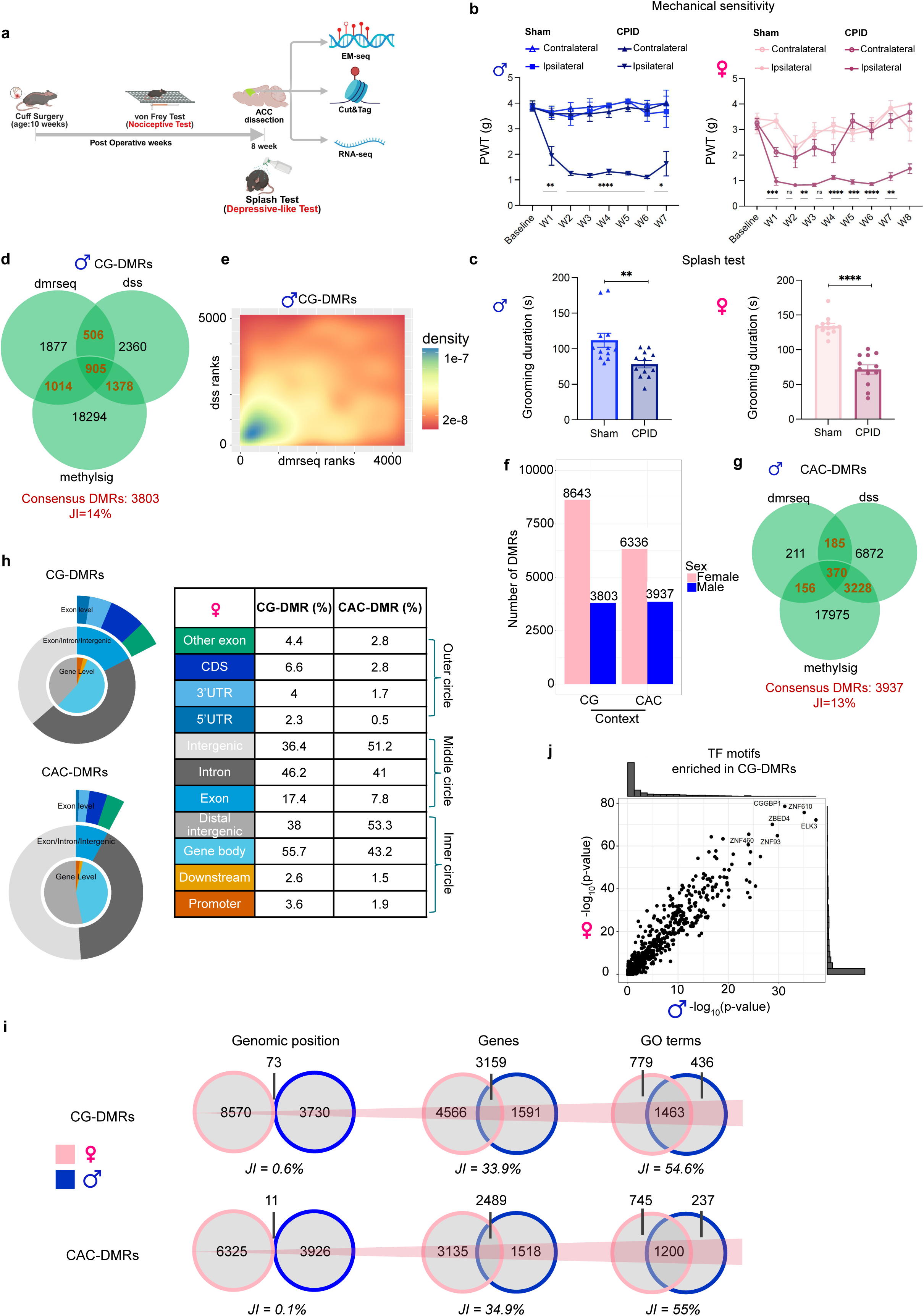
Chronic Pain-Induced Depression triggers distinct DNA methylation changes in male and female mice. **a**. Experimental design **b.** Peripheral nerve injury induced an ipsilateral mechanical hypersensitivity in both male and female mice. **c.** At 8 weeks post-surgery, male and female mice with nerve injury also displayed depressive-like behaviors, as shown by decreased grooming duration in the Splash test. **d.** Venn diagram of differentially methylated regions identified in males in the CG context (CG-DMRs) using 3 algorithms: dmrseq, methylsig, and DSS. Fourteen percent of DMRs were identified by at least 2 algorithms (JI, Jaccard Index). **e.** Density plot of ranks of DMR identified in common by DSS and dmrseq (in males, in the CG context). **f.** Number of DMR identified in each cytosine context (CG and CAC), and in each sex. **g.** Venn diagram of DMRs identified in males in the CAC context (CAC-DMRs) using the same 3 algorithms and parameters as in the CG context (see Methods). **h**. Distribution of CG- and CAC-DMR along gene features, in females. **i.** Convergence of chronic pain-induced DNA methylation changes, depicted in the CG (top) or CAC (bottom) contexts: overlaps between females and males are shown for genomic positions of DMRs (left), associated genes (middle), and related Gene Ontology (GO) terms (right). **j.** Scatter plot of enrichments of Transcription Factor (TF) binding motifs in female CG-DMRs compared to male CG-DMRs (Spearman correlation, r=0.96, p<1e-300).

To measure DNA methylation, we prioritized a method based on enzymatic conversion (EM-Seq) that improves coverage of GC-rich regions compared to the previous gold-standard of bisulfite conversion [47] (n=6/group/sex, n=24 total; Fig.S1a-b). Sequencing data were generated at an average per-sample coverage of CG sites of 9.3±0.004, and used for the identification of differentially methylated regions (DMR) in each sex. Because a consensus has yet to be reached for DMR calling [48], we combined 3 algorithms: methylsig, dmrseq, and DSS. In the canonical CG context, this led to: 21661 and 22154 DMRs identified using MethylSig in males and females, respectively; 4302 and 8010 DMRs with dmrseq; 5153 and 17967 DMRs using DSS (TableS1). Consistent with our recent work in a mouse model of addiction [49], we observed low overlaps among these results, with Jaccard Indices (JI) of 14% or 23% when considering DMR identified by at least 2 of the 3 DMR callers in males or females, respectively (Fig.1d, S1c). Importantly, these “consensus” DMR corresponded to subsets of most significant results generated by each tool (Fig.1e, S1d). Therefore, in a conservative approach, they were prioritized for downstream analyses, resulting in 3803 and 8643 CG-DMR in males and females, respectively (Fig.1f, TableS1).

We next performed a similar DMR calling for non-CG DNA methylation, given accumulating evidence that this brain-enriched epigenetic mark contributes to psychiatric disorders [21, 50–52]. We focused on CAC sites, where non-CG DNA methylation levels were highest in our ACC data (Fig.S1b), consistent with previous studies in other brain regions [49, 51, 53–55]. The exact same tools as for the CG context were used, leading to the identification of 21759 and 18539 DMRs with MethylSig in males and females, respectively; 922 and 2019 with dmrseq; and 10685 and 21417 with DSS (TableS1). We again prioritized loci identified by at least 2 methods, resulting in 3937 (JI=13%) and 6336 (JI=18%) consensus CAC-DMR in males and females, respectively (Fig.1f-g,S1c). These CG- and CAC-DMRs were widely distributed throughout the genome, and consisted of bidirectional alterations (Fig.S1e), indicating that enzymatic pathways involved in both methylation and demethylation of the DNA were recruited. At these loci, a principal component analysis (PCA) nicely separated sham and CPID groups in each context and sex, as expected (Fig.S1f). Interestingly, DMRs were predominantly located outside promoter regions, with significant differences between the 2 cytosine contexts (χ^2^ test, p<2.2e-16). While CG-DMRs were most frequently located within gene bodies (55.7%), CAC-DMRs were mostly in distal intergenic regions (53.3%, Fig.1h), suggesting distinct forms of plasticity.

Next, we wondered whether common genomic sites were affected in both sexes, and assessed overlaps among male and female DMRs. Remarkably, only 73 (JI=0.6%) and 11 (JI=0.1%) of DMRs overlapped in the CG and CAC contexts, respectively (Fig.1i, TableS2). Although rare, these overlaps were nevertheless more frequent than expected by chance (100,000 permutations with regioneR; p=1e-5 in the CG context; Fig.S2a; *Methods*), reflecting the high number of cytosines interrogated by EM-Seq. Similarly, female DMRs observed in each cytosine context were located closer to their male counterparts than expected by chance (p=1e-5 in both contexts; Fig.S2b). Then, we explored whether CPID-induced CG and CAC DMRs were occurring at loci defined by preexisting baseline sex differences. Consensus sex-DMR were identified by comparing female and male Sham control groups, using the same tools and approach described above. Results indicated that sex-DMR (n=1641 in CG context, n=1119 in CAC context) showed little overlap with CPID-DMRs, with JI equal to 4.8% and 2.7% in the CG and CAC contexts, respectively (Fig.S2c). Therefore, CPID was associated with highly sex-specific but non-random changes in DNA methylation patterns, which were not determined by preexisting sex differences.

We next used rGREAT to annotate DMR to genes located in *cis [56]*. Compared with JI observed for genomic coordinates of DMR, overlaps among genes associated with them increased to 33.9% and 34.9% in the CG and CAC contexts, respectively (Fig.1i, TableS3). This indicated that CPID-induced epigenetic changes impacted partly similar groups of genes across sexes. We then identified the GO terms enriched for DMR-associated genes. Remarkably, JI further increased, with 54.6 and 55.0% of GO terms being common between males and females in the CG or CAC contexts, respectively. The most significantly enriched GO terms related to nervous system development and synapse organization, particularly for CAC-DMR in both sexes (TableS4). Therefore, although CPID triggered methylomic changes at distinct genomic sites across sexes, this epigenetic reprogramming converged onto progressively overlapping sets of genes and biological pathways, representing a significant functional convergence. In parallel to this convergence between sexes, another one was identified across the 2 cytosine contexts (Fig.S2d). While overlaps among CG- and CAC-DMRs were low in both males (n=18, JI=0.2%) and females (n=57, JI=0.4%), they increased for the genes (n=1938; JI=28.4%; females: n=3747, JI=39.0%) and GO terms (n=1071; JI=47.2%; females: n=1541, JI=58.2%) that were associated to those DMR. Hence, the 2 forms of plasticity occurring at CG and CAC sites affected distinct loci, but functionally converged.

Because we were surprised by the extent of these sex differences (using by definition arbitrary thresholds for the identification of DMR), we next sought for threshold-free patterns of similarity in CPID-induced DNA methylation changes. To do so, we used RedRibbon [57], a method for rank-rank hypergeometric overlap compatible with large datasets. Applied to 500-bp non-overlapping bins throughout the genome (∼4.6 million bins), RedRibbon identified a significant degree of convergence across sexes (Fig.S3a-c). In the CG context, the most significant overlap involved 121,570 bins showing increased methylation levels as a function of CPID in both females and males (-log10(pval)=44.6); in the CAC context, it was composed of 105,563 hypermethylated bins (-log10(pval)=133.9). As expected, these sex-convergent bins showed milder CPID-induced methylation differences than DMRs (Fig.S3d). They also exhibited a non-random distribution as they were approximately 10 times closer to DMRs than expected by chance (e.g. in females: 3.4 kb observed compared to 40.9 kb expected; 100,000 permutations, p=1e-5 in both sexes; Fig.S3e). Altogether, these results indicate that chronic pain induced numerous but mild methylomic changes at shared loci across sexes (identified by RedRibbon), accompanied by fewer but stronger differences at sex-specific sites (DMRs).

We then characterized the transcription factors (TFs) potentially modulated at DMRs (TableS5). To do so, we used TFmotifView to assess the enrichment of TF binding motifs [58]. Interestingly, although CG-DMRs were located at distinct loci in each sex (Fig.1i), they nevertheless exhibited enrichment for strikingly similar binding motifs (CG context: p<1e-300; r=0.96; Fig.1j). This was illustrated by most significantly enriched TFs (Fig.S4a-b), as 5 out of the top 10 were shared between females and males CG-DMRs (ELK3, ZNF610, CGGBP1, ZNF93 and ZBED4). Some of these TFs have already been studied for their roles in chronic pain or synaptic plasticity (e.g., ARNT [59], NEUROD2 [60]), while others represent candidates for future investigation. Of note, no such correlation across sexes was observed in the CAC context (r=-0.03, p=0.37; Fig.S4c), at least when considering the full sets of DMRs (see below for the analysis of subgroups of DMRs defined using histone data). Overall, we conclude that CPID-induced DNA methylation changes occurred at distinct loci in each sex but modulated TFs that were partly similar across sexes.

### Analysis of DNA methylomic features confirms sex differences in CPID-induced epigenetic plasticity

To deepen the characterization of the methylomic impact of CPID, we implemented 2 additional strategies. The first used methylSeekR [61] to identify 2 types of methylomic features (Fig.2a) and their genomic localization: CG-rich unmethylated regions (UMRs) and CG-poor lowly methylated regions [62] (LMRs). As expected, UMRs primarily overlapped with promoters (∼69%), while LMRs mainly overlapped with gene bodies (54%) and distal intergenic regions (∼39%; Fig.2b), reflecting the established notion that UMRs and LMRs correspond to CG islands and distal regulatory elements, respectively [62]. As additional validation, we confirmed the previously described enrichment of these features for 2 histone modifications, H3K4me2 (using external data [63]) and H3K4me1 (our data; see below). Visual inspection revealed patterns consistent with expectations, which were confirmed by genome-wide quantification (Fig.S5a-b). Next, we identified successive CG sites that either gained or lost UMR/LMR status as a function of chronic pain, defined as diffLMRs and diffUMRs (see *Methods* and Fig.2c). This resulted in 276 and 169 diffUMRs in females and males, as well as 1051 and 541 diffLMRs in females and males, respectively (TableS6). Unlike DMRs, which exhibited high and relatively homogenous methylation differences among Sham and CPID groups, diffLMRs and diffUMRs showed lower and more variable differences (Fig.S5c). Interestingly, although these regions poorly overlapped with DMRs, they nevertheless provided a clear separation of groups (PCA; Fig.S5d), revealing a complementary biological signal. Next, we observed that, similar to DMR, diffLMRs from males and females functionally converged: their overlap in terms of genomic position was limited (diffLMRs JI=0.82% and diffUMRs JI=0.90%), but gradually increased at the gene (diffLMRs JI=9.9% and diffUMRs JI=4.2%) and GO levels (diffLMRs JI=30.9% and diffUMRs JI=11.9%) (Fig.2d, S5e, S6a; TableS3,4,6).

**Figure 2.**
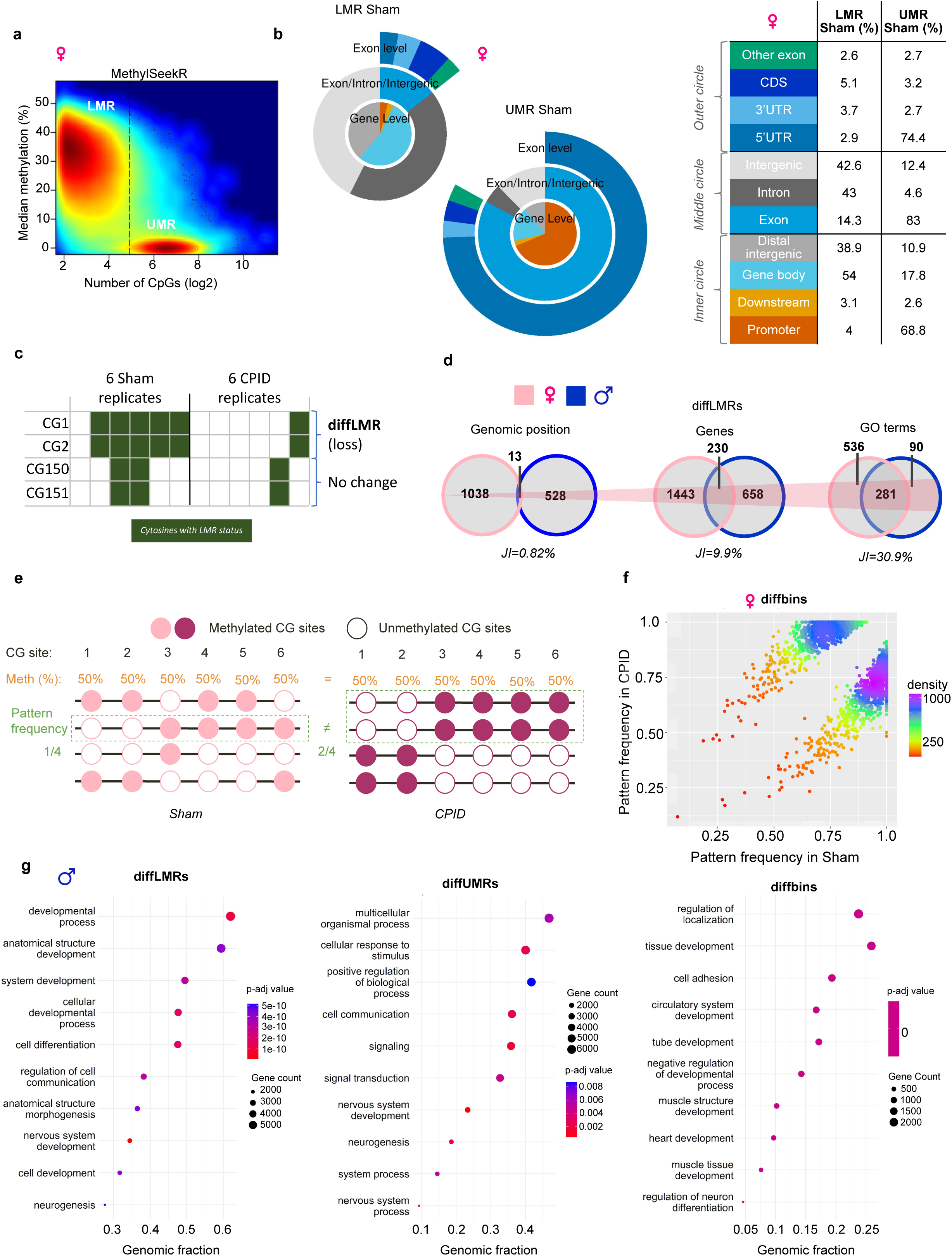
Chronic Pain-Induced Depression induces sex-specific changes in regulatory elements and single-read methylation patterns that converge on similar genes and biological pathways. **a.** Identification of female unmethylated regions (UMRs) and lowly methylated regions (LMRs) using MethylSeekR [61]. **b.** Distribution of UMR and LMR along gene features, in females. **c.** Approach for the identification of successive CG sites showing chronic pain-induced switches in LMR (diffLMRs) or UMR (diffUMRs) status. **d.** Convergence of chronic pain-induced diffLMRs: overlaps between females and males are shown for genomic positions (left), associated genes (middle), and enriched GO terms (right). **e.** Approach for the identification of chronic pain-induced changes in single-read methylation patterns (diffbins; see Methods). **f.** Scatter plot of the frequencies of single-read methylation patterns observed at diffbins in CPID (y-axis) and Sham (x-axis) groups. **g.** Top ten GO terms enriched in male diffLMRs, diffUMRs and diffbins.

The second strategy consisted in the analysis of single-read methylation patterns, using CluBCpG [64, 65]. This algorithm is designed to identify binary methylation states of successive CG sites along individual sequencing reads (i.e., single alleles), with the goal of unravelling differences that may remain masked when methylation levels are averaged across reads, such as in DMR calling (Fig.2e). We used this approach to analyze methylation patterns in non-overlapping 150-bp genomic bins, and then identified bins that showed significant changes in pattern frequency between Sham and CPID groups, defined as diffbins (*Methods*; Fig.2f). This resulted in the identification of 2254 and 4877 diffbins in females and males, respectively (TableS6), which again supported a clear segregation of CPID and Sham groups (Fig.S6b). Interestingly, these diffBins: (i) showed limited overlaps with DMR or diffUMR/LMR (Fig.S6c-e), consistent with our hypothesis that ‘epipolymorphisms’ unveil a distinct signal, and (ii) again corresponded to sex-specific loci that converged at gene and GO levels (Fig.2g, S5e, S6a, TableS3,4). Overall, results from these 2 strategies reinforce the principle of a DNA methylation sex divergence with functional convergence.

### CPID-induced changes in histone landscapes

Given the surprisingly strong sex differences observed for DNA methylation, we next sought to generalize our findings to other epigenetic processes, focusing on histone modifications. We used Cut&Tag [66] to analyze 2 activating marks primarily associated with enhancers and promoters, H3K27ac [67, 68] and H3K4me1 [69, 70], as well as a repressive mark, H3K27me3 [71–74] (n=6 libraries/group/sex/histone mark; n=72 libraries total). After peak calling, we confirmed that: (i) these data clustered first according to the repressing/activating function of each mark, and then by sex (Fig.3a); (ii) each mark displayed the expected positive and negative correlation with gene expression (Fig.3b, S7a); and (iii) H3K27ac and H3K4me1 were enriched in gene bodies (Fig.3c, S7b), particularly at Transcription Start Sites (TSS), consistent with previous reports [21, 75, 76].

**Figure 3.**
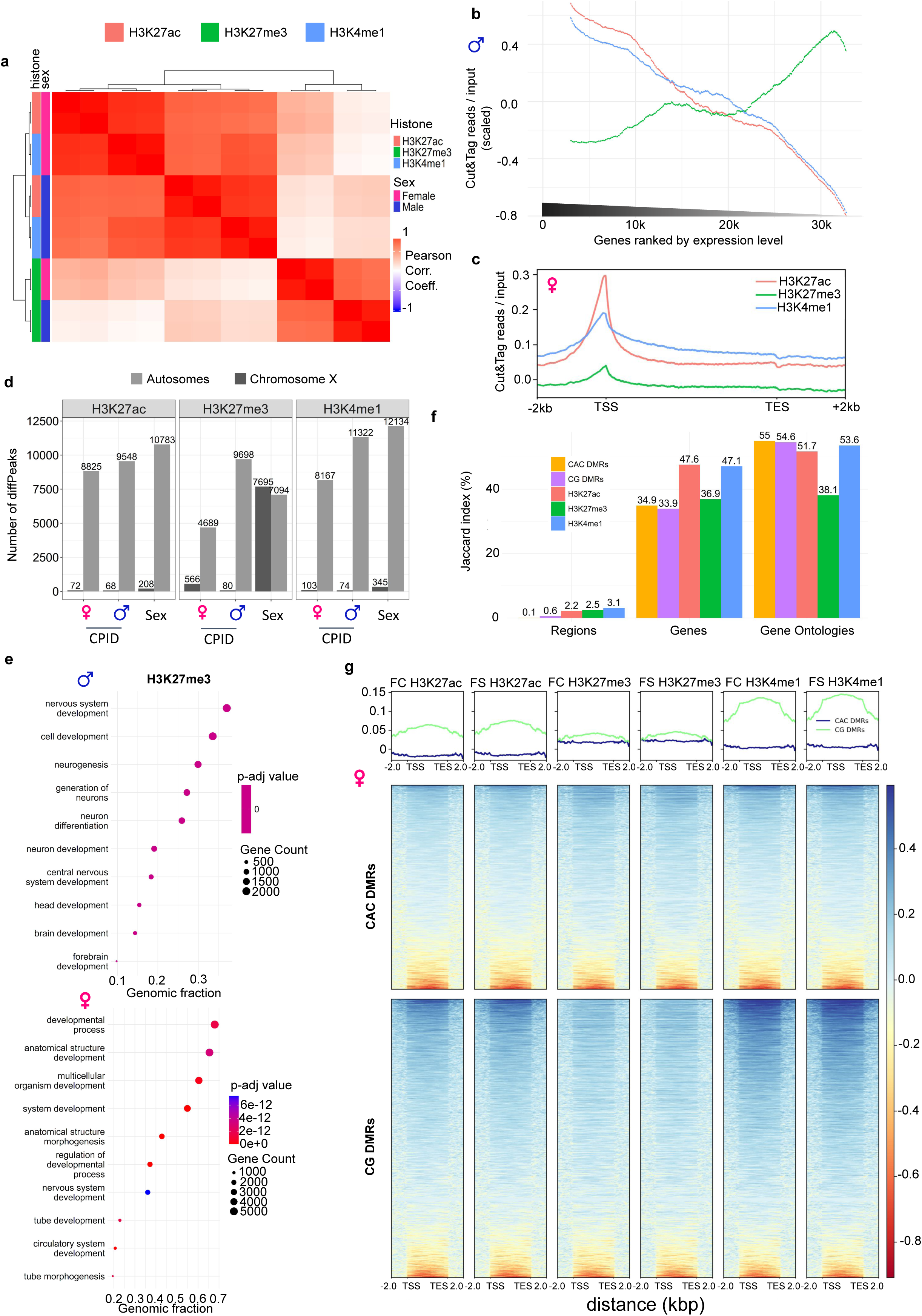
Chronic Pain-Induced Depression recruits sex-specific changes in histone modifications that show functional convergence. **a.** Hierarchical clustering of histone data (libraries were averaged among each histone mark and sex). **b.** Abundance of histone marks according to gene expression, in males (see Fig.S7a for female data). **c.** Gene profiles of histone marks, in females (see Fig.S7b for male data). **d.** Number of differential peaks (diffPeaks) identified as a function of sex or chronic pain. **e.** Top ten GO terms enriched for H3K27me3 diffPeaks in males and females. **f.** Convergence observed between males and females, for each histone mark, in terms of genomic positions, associated genes and GO terms (compared to previous results for CG- and CAC-DMRs). **g.** Heatmaps of histone marks at female CG- and CAC-DMRs (FC, Female CPID; FS, Female Sham).

Following these validations, we performed a differential analysis to identify peaks where the abundance of each mark was altered by sex or CPID (defined as diffPeaks). On the X chromosome, sex differences were more pronounced for H3K27me3 than for the other 2 marks (Fig.3d), reflecting the well-characterized and high abundance of this mark on the inactive X chromosome of females [77]. In contrast, sex differences for H3K27ac and H3K4me1 were primarily located on autosomes. Regarding the impact of CPID, we first noted that it was strong enough to be detectable by PCA (using all peaks, regardless of significance; Fig.S7c-h). Second, similar proportions of diffPeaks were identified in each sex: 8897 and 9616 for H3K27ac in females and males, respectively; 5255 and 9778 for H3K27me3; 8270 and 11396 for H3K4me1 (Fig.3d, S8a, TableS7). Biological pathways enriched for these changes included synapse organization and nervous system development, consistent with our previous DMR results (Fig.3e). Importantly, although these diffPeaks were located at sex-specific loci, they progressively converged at the level of associated genes and GO terms (Fig.3f, Fig.S8b, TableS8,9), similar to our previous DNA methylation results (Fig.3f), thereby reinforcing the principle of epigenetic divergence.

Next, to explore how this histone dataset could deepen our understanding of CPID effects on DNA methylation, we plotted histone profiles along the 2 types of CG- and CAC-DMRs. A stronger enrichment of H3K27ac and H3K4me1 was observed at CG-DMRs compared with their CAC counterparts (see Fig.3g for females), indicating that the former occurred more frequently within euchromatic or enhancer regions [78]. Interestingly, a similar pattern was observed in males (Fig.S8c), suggesting that it may reflect distinct mechanistic cross-talk between histones and the 2 types of DNA methylation. We further noticed substantial heterogeneity within each DMR type, which prompted us to apply an unsupervised k-means clustering approach. This identified clusters of DMRs that were similar across both cytosine contexts (Fig.4a-b, and TableS10). The first cluster (C1) was enriched for H3K27me3 and corresponded to heterochromatic regions, whereas the second (C2) was characterized by high levels of H3K27ac and H3K4me1, consistent with active enhancers [79]. The third cluster (C3) exhibited lower levels of all 3 histone modifications and likely represented poised enhancers. The remaining clusters were depleted of all 3 marks and were not investigated further. Reflecting these patterns, genes associated with C3-DMRs displayed significantly higher expression than those associated with C1-DMRs but lower expression than those associated with C2-DMRs, in both cytosine contexts (Fig.4c). These DMR subgroups also differed in their genomic distribution relative to genes, as C2-DMRs (C2) were more frequently located in introns, consistent with their potential role as regulatory elements (Fig.4d). Importantly, the same clusters were observed in a similar analysis in males (Fig.S9a-b), and genes associated to a given cluster and sex were enriched for genes associated with that same cluster in the other sex (Fig.4e, S9c). These reproducible clusters may therefore reflect distinct forms of interaction between DNA methylation and histone marks. To support this hypothesis, we analyzed TF binding motifs enriched within each cluster of DMRs (TableS11). Interestingly, this analysis: (i) unveiled strong correlations between females and males for C1, C2 and C3 clusters of CAC-DMRs (Fig.4f, S9d-e), contrasting with our initial results for the full set of CAC-DMRs (Fig.S4c); and (ii) showed that the TFs with highest enrichment were specific to each cluster (e.g. KLF9 and PATZ1 in C2, MEF2D and POU3F1 in C3 CAC-DMRs). Overall, these results indicate that chronic pain recruited distinct classes of DMRs defined by various histone landscapes. These classes were similar across both sexes and cytosine contexts, yet targeted sex-specific loci and likely modulated distinct sets of TFs.

**Figure 4.**
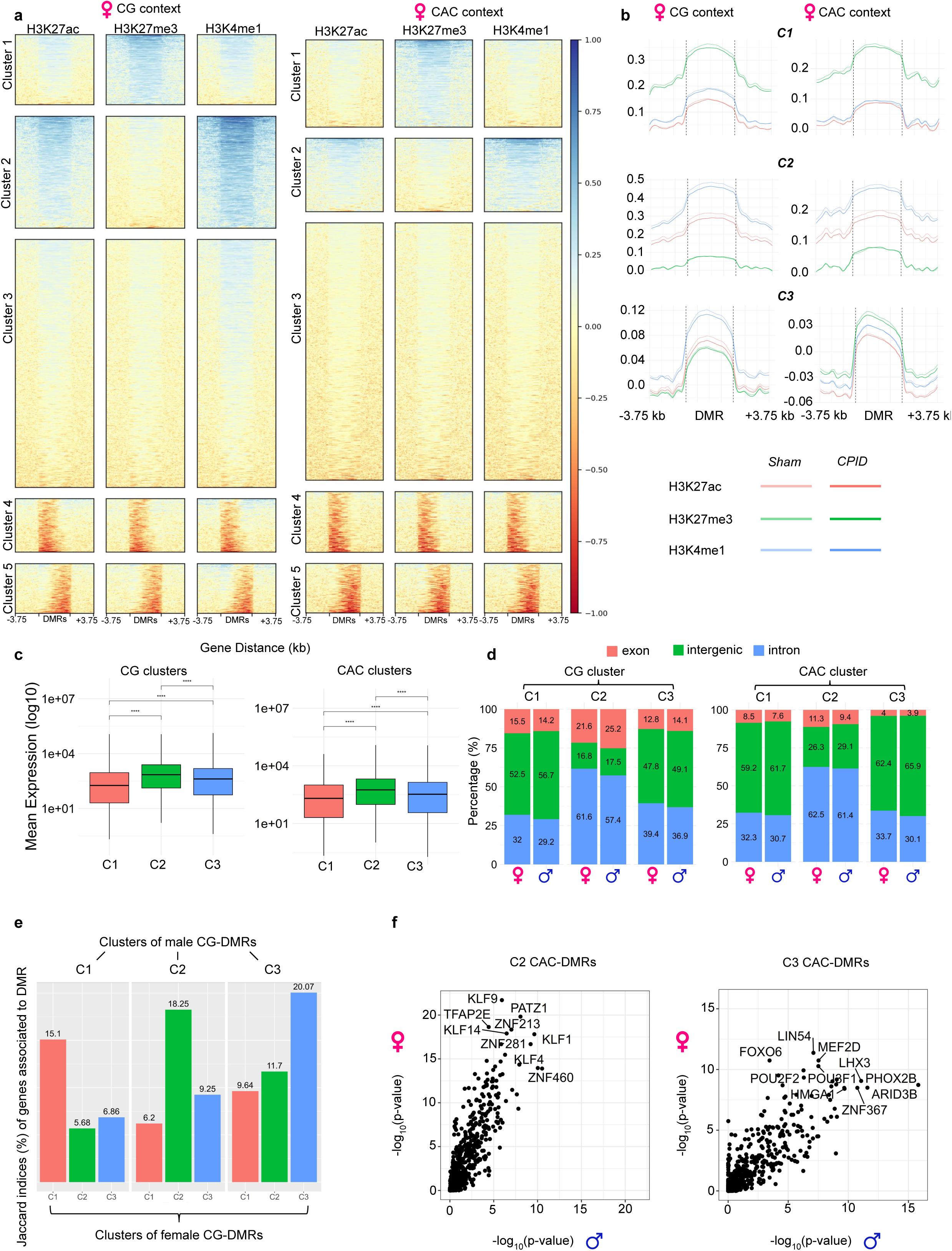
Histone marks differentiate 3 types of chronic pain-induced DNA methylation changes, which are conserved across sexes. **a.** Heatmaps of histone marks at clusters of CG- and CAC-DMRs identified in females (see Methods, Fig.S9a for female data, and TableS10). **b.** Profiles of histone marks for each cluster of female CG-DMR and CAC-DMRs. **c.** Mean expression of genes associated with each cluster of CG-DMR (p<1e-15 for all comparisons) and CAC-DMRs (p<1e-6 for all comparisons). **d.** Distribution along gene features of CG and CAC clusters of DMR, in each sex. **e.** Overlaps between females and males of genes associated with cluster-specific CG-DMRs. **f.** Correlation of transcription factor motif enrichment between female and male cluster 2-DMRs (left panel; Spearman correlation, r=0.92, p<1e-300), and between female and male cluster 3-DMRs (right panel; r=0.86, p<2.7e-297).

### Network integration identifies gene modules and biological pathways impacted by CPID at multiomic levels

Having established the multilayered effects of chronic pain on the epigenome, we next investigated the transcriptional consequences of these alterations. To this end, we performed RNA-seq on bulk ACC tissue (n=6 per Sham and CPID group in each sex; n=24 total), followed by differential expression analysis. This identified 712 and 444 differentially expressed genes (DEG; defined using DESeq2 and nominal p<0.01) in females and males, respectively (Fig.5a, TableS12), of which only 21 were shared between sexes (JI=1.8%, with 20/21 dysregulated in similar directions; Fig.5b). Consistent with this limited overlap, Gene Ontology (GO) analysis revealed only a small number of enriched terms common to both sexes (e.g., synapse organization). Most enrichments were sex-specific, including nucleosome assembly and regulation of angiogenesis in males, and axonogenesis and axon ensheathment in females (Fig.5c, TableS13). Intriguingly, the enrichment of neurovascular-related transcriptional changes in males (which was absent in females; Fig.S10a) contrasts with recent findings from depression models based on chronic social defeat or chronic variable stress, in which neurovascular dysregulation was reported in females but not males [80]. This raises the possibility that these alterations may contribute to depressive-like behaviors in a manner that depends on both sex and etiology.

**Figure 5.**
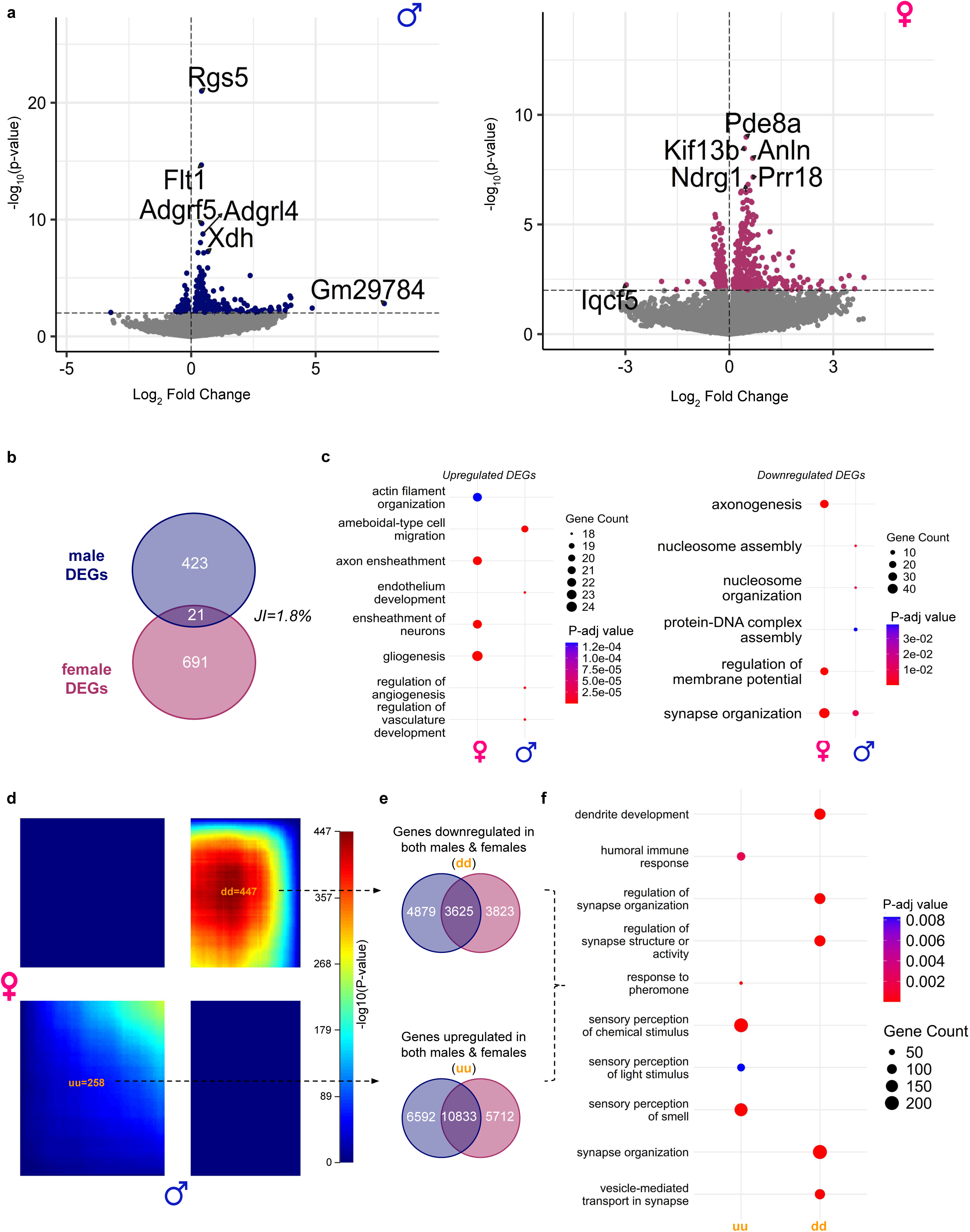
Chronic Pain-Induced Depression recruits sex-specific transcriptional signatures. **a.** Volcano plots of transcriptomic profiles in males and females, with differentially expressed genes (DEGs, p-value<0.01) shown in blue and pink, respectively. **b.** Venn diagram of DEGs identified in males and females (JI, Jaccard Index). **c.** Top GO terms (BPs) enriched for chronic pain-induced up- or downregulated DEGs in females and males. **d.** Threshold-free comparison of transcriptomic effects of chronic pain across females and males (using RRHO2 for iterative hypergeometric testing). **e.** Venn diagrams of the 2 best overlaps among genes commonly down- (d/d) or upregulated (u/u) in both females and males. **f.** Top GO terms (BPs) enriched in the d/d or u/u gene sets.

Next, similar to our DNA methylation analysis, we performed a threshold-free comparison of the male and female transcriptional signatures. This revealed a strong concordance between sexes, with large sets of genes showing consistent upregulation (uu; p=1e-258, n=10833) or downregulation (dd; p=1e-447, n=3625; Fig.5d,e, TableS14) in both males and females. Notably, these concordantly regulated genes were significantly enriched for GO terms related to synaptic processes (Fig.5f, TableS14), extending the findings from the DEG analysis and indicating that chronic pain triggered shared alterations in synapse-related transcriptional programs across sexes. Overall, these results indicate that although the majority of DEGs were sex-specific, a broader pattern of transcriptomic convergence emerged when gene expression changes were analyzed in a threshold-free manner.

Finally, to integrate our epigenomic and transcriptomic datasets, we performed gene co-expression network analysis using MEGENA (Multiscale Embedded Gene co-Expression Network Analysis; see [81]). When applied separately to male and female transcriptomes, MEGENA identified modules of co-expressed genes that showed substantial similarity, with 64% of modules identified in one sex exhibiting preserved co-expression in the other (see *Methods* and Fig.S10b). We therefore combined female and male data to construct a unified network and facilitate biological interpretation. Then, we assessed the impact of CPID on each module by testing for enrichment of gene sets identified across the different omic layers, using Fisher’s Exact Test (FET). Most correlations among enrichments observed for pairs of layers were significant (Fig.6a), even between sexes, indicating that pain-induced changes in DNA methylation and histone modifications converged on similar modules. These enrichments were then averaged, using Stouffer p-values, to generate in each sex a ranking of modules from the least to the most significantly impacted by chronic pain (Fig.6b). Results showed that these rankings were strongly correlated between females and males (r=0.97, p=4.3e-12; Fig.S10c). Strikingly, among the 8 and 9 modules impacted in females and males, respectively (Stouffer’s p<0.01, see TableS15), 8 were common to both sexes (m3, m6, m9, m10, m8, m17, m19, m12). Overall, these findings extend at the module level the notion of functional convergence arising from sex-specific epigenetic adaptations that was previously observed for each individual omic layer.

**Figure 6.**
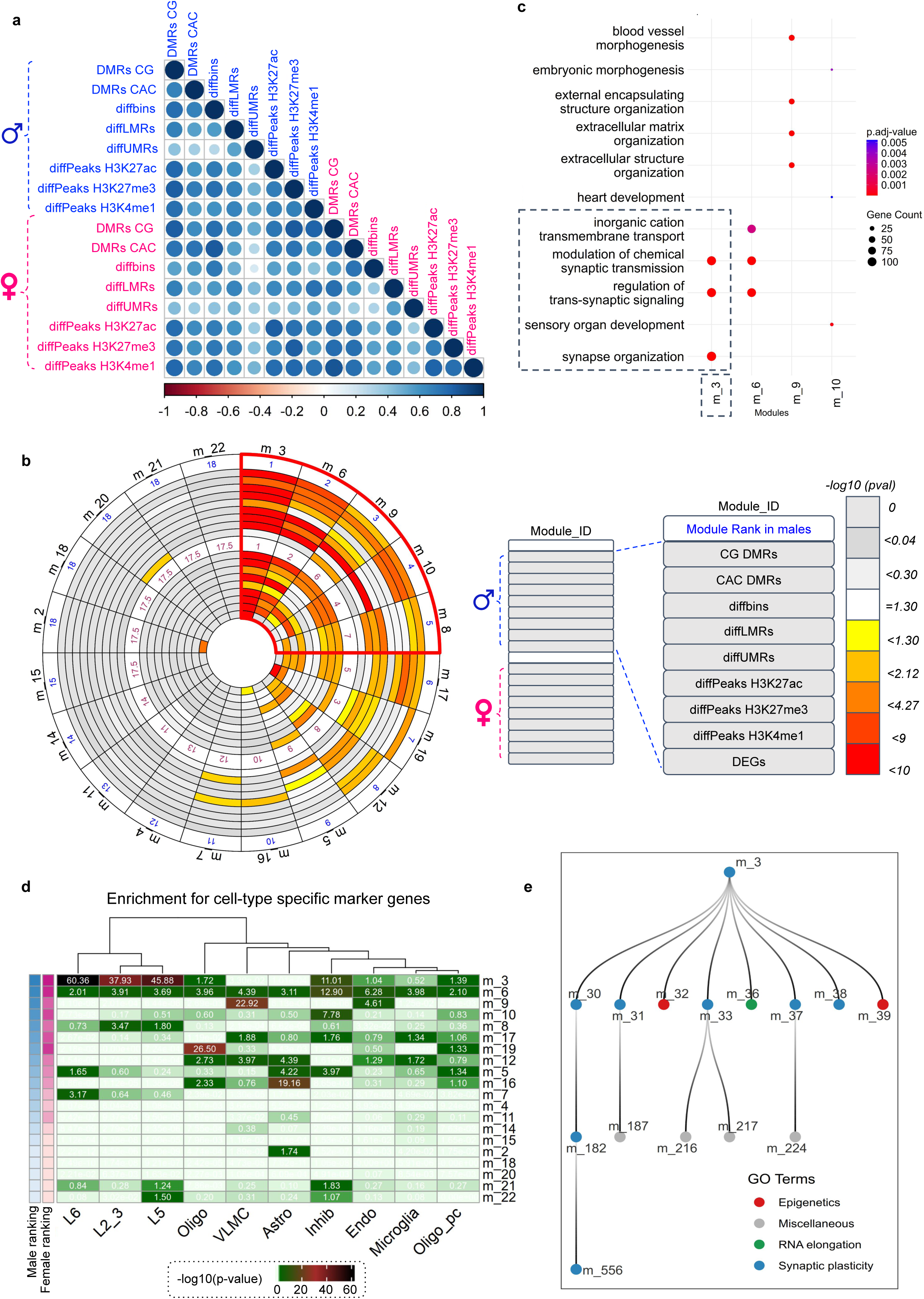
Chronic Pain-Induced Depression epigenetically reprograms similar gene modules and cell types in both sexes. **a.** Pearson correlations among enrichments of MEGENA co-expression modules for pairs of omic layers. **b.** Circos plot depicting the degree of enrichment (Fisher’s exact test), within each first-generation MEGENA module, of every type of chronic pain-induced differences at the level of DNA methylation (DMRs, diffbins, diffLMR/UMRs), histone marks (diffPeaks), or the transcriptome (DEGs). Enrichments across omic layers were averaged using Stouffer’s p-values and used to rank modules in each sex. Similar modules showed highest multiomic enrichment in males (outer circles) and females (inner circles). **c.** Top GO terms (BPs) enriched in the 4 MEGENA modules most strongly enriched for chronic pain-induced multiomic effects. **d.** Heatmap of the enrichment of MEGENA modules (shown as rows, ranked using Stouffer p-values in males) for gene sets defining cortical cell-types (columns, using data from [82]). Abbreviations: L2_3, L_6, L_5, pyramidal neurons from the corresponding cortical layers; VLMC, Vascular and LeptoMeningeal Cells; Oligo, oligodendrocytes; Oligo_pc, oligodendrocyte progenitor cells. **e.** Child modules of the m3 parent module (see main text for details).

We then characterized the biological functions of modules most significantly affected by chronic pain at multiomic levels. GO terms enriched in these modules were related to the synapse for m3 and m6, to the endothelial system for m9, to developmental processes for m10, among others (Fig.6c and TableS15 for full results). To associate these alterations with cortical cell types, we used 3 independent external single-cell data that were generated in the mouse prefrontal cortex, and defined marker genes [82–84] (Fig.6d, S11a-b, TableS16). The 3 datasets consistently indicated that most affected modules were enriched in marker genes of excitatory (m3, m8) and inhibitory neurons (m6, m10), as well as endothelial cells (m9, consistent with the GO enrichment observed for that module), while those enriched in marker genes for oligodendrocytes (m19) or astrocytes (m16) were comparatively less impacted. Finally, we considered the hierarchical organization of MEGENA networks, which identifies parent-child relationships among modules. Interestingly, the successive generations of children modules originating from the most significantly affected parent one, m3, showed multiple enrichments for GO terms related to epigenetic processes and synaptic functions. For example, these included ‘histone binding’ and ’chromatin binding’ (enriched in m32); ‘histone-modifying activity’ and ’histone acetyltransferase activity’ (m39); as well as ‘action potential’ (m30, Biological Process, BP) or ‘postsynaptic specialization membrane’ (m31) (Fig.6e, TableS15). Overall, these results indicate that although they have been relatively understudied in that context [24], epigenetic processes may represent important mediators of the pain-depression comorbidity that are particularly impaired in excitatory and inhibitory ACC neurons. Studying how these epigenetic changes mediate the synaptic dysfunction that we repeatedly identified across multiple omic layers therefore represents an appealing perspective for future work.

## Discussion

Chronic pain-induced depression involves alterations affecting the ACC, a key brain region for attributing emotional valence to nociceptive information. Although both male and female mice develop depressive-like behaviors after the induction of chronic neuropathic pain, whether sex-specific molecular mechanisms are implicated remains largely unexplored [18, 85, 86]. In this context, the present study sought to characterize sex differences in the epigenomic and transcriptomic plasticity recruited in the ACC in our CPID model.

Pain-depression comorbidity has been shown to rely at least partly on an imbalance between neuronal excitation and inhibition (E/I), leading to a shift toward cortical hyperactivity [87]. Within the ACC, several studies have reported increased c-Fos expression and enhanced intrinsic excitability and synaptic activity of pyramidal neurons, consistent with elevated excitatory drive [43, 87]. In parallel, a reduction in inhibitory control has also been described, including decreased inhibitory synaptic transmission [87]. The present study uncovers consistent multiomic changes that likely provide the molecular scaffold for this electrophysiological model. Specifically, we found that pain-induced epigenetic changes were enriched in highly inter-connected gene modules, which were strongly associated with synaptic plasticity, excitatory and inhibitory ACC neurons, as well as epigenetic regulators. We therefore propose that these epigenetic changes contribute to the dysregulation of excitatory and inhibitory circuits, persistent E/I imbalance and ACC hyperactivity observed in chronic pain [88].

A main originality of our work lies in the integrative analysis of multiple epigenetic layers that include DNA methylation and histone modifications. Although the relationships among these layers are well-characterized in physiological contexts, their cross-talk during chronic pain had not been previously characterized [24, 89, 90]. In addition, while previous studies focused on the sole CG context or a limited fraction of the genome [27–35, 37, 38], we examined both CG and CAC methylation, in the full genome. Our results demonstrate that both types of DNA methylation were substantially remodeled by chronic pain, with striking differences. CG- and CAC-DMRs showed minimal overlap, distinct distributions relative to genes, and different associations with transcription factors, suggesting that the two types of adaptations were embedded within distinct regulatory environments. Nevertheless, both CG- and CAC-DMRs grouped into 3 clusters characterized by shared histone profiles. They also converged on common gene co-expression modules and biological pathways. These findings therefore support a model in which CG and non-CG methylation constitute complementary yet mechanistically distinct layers of regulation [21, 91] that act in parallel to shape the transcriptional adaptations associated with chronic pain.

The most significant contribution of this study concerns the influence of sex on the epigenetic plasticity involved in chronic pain-induced depression. Sex differences have been documented across multiple levels of brain organization, including gene expression, cellular composition, circuit architecture, and behavior [92–94]. Despite growing recognition that both chronic pain and depression differ markedly between males and females in prevalence, symptomatology, and treatment response, the molecular mechanisms underlying these differences remain incompletely understood. In particular, no previous preclinical studies have examined multiple epigenetic layers simultaneously in both sexes at genome-wide resolution. Our analyses revealed extensive sex differences in the epigenomic response to chronic pain. Across both DNA methylation and 3 histone marks, the loci affected by CPID showed remarkably limited overlap between males and females. These changes were distributed throughout the genome and were largely distinct from baseline sex differences, which were predominantly concentrated on the X chromosome. Thus, the sex-specific molecular response to chronic pain cannot be readily predicted from constitutive epigenetic differences between males and females. Despite this pronounced divergence at the level of individual loci, several observations pointed to a higher degree of functional convergence. In both sexes, CPID-associated epigenetic alterations displayed similar genomic organization, recruited related transcriptional regulatory programs, and ultimately converged on overlapping gene co-expression modules and biological pathways. These findings suggest that males and females engage similar epigenetic mechanisms that act at distinct loci to achieve comparable transcriptional and functional outcomes. More broadly, they raise the question of the mechanisms that may be responsible for generating such divergent epigenomic responses to chronic pain in males and females. Future studies will be needed to determine how this divergence may be’ shaped by biological factors that include fluctuations in gonadal hormones [39], sex-dependent features of chromatin architecture [95], as well as sexually dimorphic epigenetic regulators [96] (such as the X-linked histone demethylases, Kdm5c and Kdm6a, and their Y-linked counterparts, Kdm5d and Uty).

This study has limitations. The first is that we performed our epigenetic investigations when chronic pain was associated with depressive-like behaviors. Our CPID model is based on a long-lasting pain experience that manifests by an initial mechanical allodynia, followed by a period during which it is accompanied by emotional dysregulation, thereby providing a temporal dissociation between the 2 phenomena [40]. Our analyses focused on this second time-point. It is therefore possible that, in addition to mediating the emotional consequences of pain, the molecular changes that we uncovered may at least in part represent long-term consequences of pain itself. More work will be necessary to define these kinetics among the intricate consequences of the pain experience and underlying epigenetic reprogramming. The second limitation is related to cellular resolution. While spanning multiple omic layers, our analyses were conducted on bulk ACC tissue and thus did not resolve the molecular heterogeneity of individual cell populations. Consequently, adaptations affecting minor cell types may have been overlooked. In addition, our integrative approach suggested that pain-induced epigenetic changes targeted shared cellular populations in both sexes, yet it remains possible that some of these adaptations arose from partly distinct cell types in males and females. Addressing this limitation will require applying single-cell approaches—already used to study transcriptional mechanisms of pain [97, 98]—to dissect epigenomic profiles in the context of pain-depression comorbidity.

In conclusion, this study establishes the extensive and sex-specific epigenetic and transcriptomic adaptations that occur when chronic pain triggers depressive-like behaviors. Despite their divergence at the molecular level, these adaptations converge on shared functional pathways in both sexes, highlighting the potential to develop sex-specific epigenetic strategies targeting common pathophysiological processes.

## Data availability

EM-seq, CUT&Tag and RNA-seq raw and processed data have been deposited at GEO under accession numbers GSE310867, GSE310866 and GSE310870, respectively, and will be made publicly available upon publication. Any additional information required to reanalyze the data reported in this paper is available from the lead contact upon request.

## Code availability

Code for bioinformatic analyses is available on GitHub: https://github.com/pelutzlab/cpid_dnam.

## Author contributions

VVS, BA, HE, KA, NW, and IY performed experiments involving the chronic pain model, conducted behavioral testing, and processed cohorts for EM-seq and RNA-seq analysis. MB and HS performed Cut&Tag experiments. Genome Quebec did EM-seq and RNA-seq sequencing. Cut&Tag sequencing was done by IGBMC. MG led all bioinformatic analyses. ACR, YH-A and SLG contributed to part of the bioinformatics analyses. PEL designed and conducted part of the analyses. IY and PEL secured financial support. BL, IY and PEL supervised the project. MG and PEL prepared the manuscript, which all authors approved.

## Acknowledgments

We thank the Chronobiotron UAR3415 for animal care. We also thank Andre Moreira Pessoni, Arturo Marroquín-Rivera, and Samaneh Mansouri for helpful discussions about bioinformatic analyses, as well as Sarah Journée, Quentin Leboulleux, Clementine Fillinger and Robin Waegaert for their help with behavioral experiments. We made figures using Biorender.

