## Supplementary Material for "Sex differences in epigenetic mechanisms of chronic pain-induced depression"

### Content

1. Supplementary material and methods
2. Legends of supplementary figures
3. Legends of supplementary tables

### 1. Supplementary material and methods

**Nociceptive behaviors.** Mechanical thresholds were assessed using von Frey filaments, before surgery (baseline) and on a weekly basis after surgery, in order to assess neuropathy-induced hypersensitivity. During each session, mice were habituated (5 min) in individual transparent, bottomless plastic boxes placed on a mesh platform. For the testing, filaments of different ascending pressure (0.4 to 8.0 g; Bioseb, Chaville, France) were applied to the ventral surface of each hindpaw. A positive response for a given pressure corresponds to withdrawal, licking, guarding or flinching of the stimulated hindpaw for 3 out of 5 applications. The mechanical sensitivity threshold was defined as the lower filament of the 2 consecutive filaments that elicited a response.

**Splash test.** This test was performed individually on mice placed in a new cage where their dorsal coat was sprayed with a 20% sucrose solution (Erstein, France) and allowed to move freely. Total grooming time over a period of five minutes was manually recorded.

**DNA and RNA Extraction.** Twenty-four hours after the last behavioral test, mice were euthanized by cervical dislocation. The whole brain was excised and sectioned in a brain matrix (Electron Microscopy Sciences, Rodent Brain Matrices, Cat. #69022). The ACC was dissected and the tissue stored at -80°C. DNA and RNA were then co-extracted using the NORGEN RNA/DNA Purification Micro Kit (Ref. #SKU48700). All centrifugation steps were performed at 4 °C, with 10 µL of β-mercaptoethanol added to each 1 mL of SKP buffer. Each sample was homogenized in 300 µL of SKP buffer 5. The homogenate was centrifuged at 200*g for 2 minutes. For genomic DNA, the supernatant was loaded to a gDNA Purification Micro Column assembled with a collection tube, and centrifuged at 5,200*g for 2 minutes. The flowthrough containing RNA was retained on ice for RNA purification. The gDNA column was washed twice with 500 µL of Wash Solution A, each time centrifuged at 3,500*g for 1 minute, with an additional spin at 14,000*g (2 minutes) for drying. The column was transferred to a fresh 1.7 mL elution tube, and 25 µL of Elution Buffer F added. After incubation at room temperature (2 minutes), centrifugation was performed at 200*g (2 minutes) and then 14,000*g (1 minute). For RNA, 60 µL of 100% ethanol was added to every 100 µL of flowthrough, mixed by vortexing, applied to a RNA Purification Micro Column assembled with a collection tube, and centrifuged at 3,500*g (1 minute). The flowthrough was discarded and the column washed three times with 400 µL of Wash Solution A, each followed by centrifugation at 3,500*g (1 minute) and then 14,000*g (2 minutes) for drying. The column was placed in a fresh 1.7 mL elution tube, and 25 µL of Elution Solution A added, followed by centrifugation (200*g, 2 minutes, and 14,000*g, 1 minute).

**EM-seq library preparation.** gDNA was quantified using the Quant-iT™ PicoGreen® dsDNA Assay Kit (Life Technologies) and its integrity assessed on agarose gels. Libraries were generated using the NEBNext® Enzymatic Methyl-seq Kit (New England BioLabs) at the McGill University and Genome Quebec Innovation Center. With this kit, the methylation status of cytosines is revealed using an enzymatic rather than chemical (bisulfite) method of conversion [1]. Briefly, we followed the manufacturer's recommendations. DNA was fragmented and used for adapter ligation and clean-up, followed by a first enzymatic reaction using the Tet Eleven Translocation Enzyme TET2 as well as an oxidation enhancer (1 hour, 37°c). After stopping this reaction, and another clean-up, a second enzymatic reaction was performed using the APOBEC enzyme (apolipoprotein B mRNA editing enzyme, catalytic polypeptide) for deamination of the DNA (3 hours, 37°c), followed by clean-up, PCR amplification, and clean-up steps. Adapters were purchased from NEB. Libraries were quantified using the Kapa Illumina GA with Revised Primers-SYBR Fast Universal kit (Kapa Biosystems) and average size fragment determined using a LabChip GX (PerkinElmer) instrument. The libraries were normalized, pooled, denatured in 0.02N NaOH and neutralized using the HT1 buffer. The pool was loaded at 200pM on a Illumina NovaSeq S4 lane, in 150 cycles paired-end mode. A PhiX library was mixed with libraries at 5% level. Program BCL Convert 4.2.4 was used to demultiplex samples and generate FASTQ reads. The average sequencing depth was 187.6 ± 8.6 million reads/library (mean+/-sem; TableS1).

**EM-seq data processing and identification of differentially methylated regions (DMRs).** Trim galore (0.6.4) [2] was used for trimming. Trimmed reads were aligned against the mm10 reference genome using Bismark [3], and unconverted or duplicated reads discarded. As quality control, the efficiency of enzymatic conversion was examined using unmethylated Lambda spiked-in DNA. All samples showed an efficiency >99% (Fig.S1a), and were kept for downstream analyses. The methylation extraction function from Bismark (with the –cytosine_report and –CX options) was used to obtain DNA methylation data for individual cytosines. These matrices were imported as a single bsseq object in each sex, using the R (4.1.2) package bsseq (1.30.0). DMRs in the CG context were called in each sex by comparing CPID and sham samples using 3 R packages: methylsig (1.6.0), dmrseq (1.14.0) and dss (2.40.0). For dmrseq, candidate regions are defined by a methylation difference greater than 5%. Then, 10 permutations (with set.seed(123)) were used to generate the null model, and a 0.1 p-value threshold used to define DMRs. For DSS, using the 5% p-value threshold, Differential Methylated Loci (DML) were defined, and regions were defined as DMRs if they contain >= 5 CG sites and >= 5% mean methylation differences. For Methylsig, we defined a window size 500-bp and filtered for regions with a non-null coverage in all samples. The beta-binomial model was then used to calculate the differential methylation statistics, with a p-value threshold of 0.01. The same analyses were carried out for the CAC context (largest non-CG context in DNA methylation; Fig.S1b), except that DMR-calling by DSS was carried out in three batches of chromosomes (chr1-6, chr7-13, and chr14-M), to avoid the computational load on the services. Genes associated with DMRs [4] were obtained using rGREAT (2.0.2) [5] with the default "basal plus extension" parameters for association with genes (basal: 5 and 1 kb up- and downstream the TSS; extension: to the nearest gene's basal domain, extendable up to 1000 kb). The Gene Ontology (GO) enriching DMRs were inferred using rGREAT (2.0.2) [5] with the whole genome of mm10 species as background for CG and CAC DMRs, as they are widespread in the genome [6–8]. The significant BPs (padj<=0.05) were encapsulated through the RRVGO (1.10.0) package [9]. The genes and GO: Biological Processes (BP) obtained from the above method were used for assessing the functional convergence across the sexes. Genomic overlaps and meandistance significance between the DMRs were computed by the "numOverlaps" and "meanDistance" arguments in the "permTest" function using the regioneR (1.26.0) package [10]. Further, we looked for the enrichment of Transcription factor binding motifs for the DMRs from both contexts and sexes by using the TFmotifView [11].

**Identification of lowly and unmethylated regions.** DNA methylation data in the CG context were used to identify the fully-, lowly- and un-methylated regions (FMRs, LMRs and UMRs) using the R package methylseekR (1.38.0) [12]. The default methylation threshold (50%) was used for segmentation, and permutation testing indicated that n=4 consecutive CG sites were required for LMR-calling at a FDR=<5%. To validate the LMRs and UMRs, we used internal H3K4me1 and external H3K4me2 [13] histone marks and visualized them in Integrated Genomics Viewer (IGV) [14]. Following the sample-specific segmentation and validation, using an in-house script, we investigated changes in the UMR/LMR localization as a function of CPID-effect independently in each sex. To do so, differential LMRs (diffLMRs) were defined as regions containing at least 5 CG sites that individually switched from the LMR state to another (FMR or UMR) in at least 4/6 samples from one group to another (Sham/CPID). Similar criteria were used to define differential UMRs (diffUMRs; hence, although UMRs contain a minimum of 30 CG sites, diffUMRs can have fewer). The associated genes and GOs were from rGREAT (2.0.2), with a background of LMRs and UMRs.

**Identification of single-read DNA methylation patterns.** To identify the read-level methylation patterns, we used CluBCpG (0.2.5) [15], with three steps: (1) Pre-processing of CG bam files: Sort the bam files coordinates, index them, and extract the chromosomes from the resulting bam files using the samtools (1.15.1); the resulting chromosome bam files were sorted (by names) and deduplicated using the Picard (2.23.3) [16]. (2) clubcpg-coverage: Identify the reads fully covering bin sizes of 150-bp, bins aligned against >= 3 reads, and containing >=2 CGs were kept. (3) clubcpg-cluster: Group the reads in clusters presenting the same single-read methylation pattern. Here, "noise filtering" was set to false. At the same time, read depth and minimum number of reads per cluster were set to 1 to measure the frequency of single-read methylation patterns among all available reads, and to eventually identify frequency changes induced by CPID-effect. Then, independently in each sex, we used in-house scripts to select methylation patterns that were reproducibly detected (non-null coverage) in at least 4 out of 6 samples in each group (CPID and Sham). Frequencies across these two groups were compared using the Wilcoxon test, which is defined by p-value<0.005 and are identified as the significant differential bins (diffbins). Associated genes and GO enrichment of diffbins were inferred using rGREAT (2.0.2), similar to diffLMRs/UMRs.

**Cut&Tag library preparation.** Neuropathic pain was induced in additional cohorts of male and female mice. To analyze three histone marks (H3K27ac, H3K27me3, H3K4me1), ACC tissue was pooled from three mice. Tissue pools were homogenized on dry ice using the KIMBLE Dounce tissue grinder set (Merck) and resuspended in phosphate-buffered saline solution mixed with cOmpleteTM Protease Inhibitor Cocktail (PBS-PIC, Roche). Suspension was centrifuged at 1,000 g for 5 minutes at 4°C, the supernatant was discarded and the pellet was resuspended in 1 ml lysis buffer (657.5 µl of MilliQ water, 200 µl of 50% glycerol, 50 µl of 1M HEPES with pH 7.5, 50 µl of 10% NP-40, 28 µl of 5M NaCl, 12.5 µl of 20% Triton X-100, 2 µl of 0.5M EDTA with pH 8.0, 40µl of 1X PIC). Samples were incubated at 4°C for 10 minutes on a rocker, homogenized 5 times (stroke "A") in the grinder, transferred back to plastic tubes of the size 1.5 mL, and pelleted by centrifugation at 1,000 g for 8 minutes at 4°C. PBS-PIC was used to wash the nuclei pellets, followed by gentle resuspension, and filtering of remaining pieces of tissue using 50 µm pore size CellTrics® filters (Sysmex). Every sample was split into equal proportions according to the number of marks. The obtained filtrates, consisting of cell nuclei and cellular debris, were filled with a wash buffer for Cut&Tag until a final volume of approximately 1.5 ml, and kept on ice during the following steps. Nuclei were attached to magnetic nanoparticles using Concanavalin A (ConA) Magnetic Beads and Activation Buffer (Cell Signaling), as described by [17] with several modifications. In short, ConA beads attached to nuclei were put through resuspension and incubation on a rotator at 4°C overnight in 50 µl Antibody buffer (8 µL 0.5M EDTA pH 8.0 + 6.7 µL 30% BSA + 2 mL Wash buffer, with 1 µl of the undiluted antibodies). Antibodies used in this study are H3K27Ac (Anti-Histone H3 (acetyl K27) antibody - ChIP Grade, Cat no. ab4729, Abcam), H3K27me3 (H3K27me3 Antibody, Cat no. C15410195, Diagenode), H3K4me1 (H3K4me1 Antibody, Cat no. C15410194, Diagenode) and IgG (Rabbit IgG Antibody - ChIP Grade, Cat no. C15410206, Diagenode). Then, liquid was removed with a magnetic rack, and ConA beads with nuclei were resuspended in 100 µl secondary antibody (Guinea Pig anti-Rabbit IgG, Heavy & Light Chain Antibody Preadsorbed, Cat no. ABIN101961, Antibodies online) diluted 100 times in Wash Buffer (1 mL 1M HEPES pH 7.5 + 1.5 mL 5M NaCl + 12.5 µL 2M spermidine and fulfilled to 50 mL with MilliQwater and one diluted tablet of protease inhibitor). On a rotator, samples were incubated at room temperature for 1 hour. After the removal of liquid and washes with Wash Buffer for three times using a magnetic rack, recombinant pA-Tn5 protein (pA-Tn5 Transposase loaded from Diagenode, Cat no. C01070001-30) was bound to the current antibody-protein complex (100 µl of diluted pA-Tn5 which was diluted for 250 times in DIG-300 (1 mL 1 M HEPES pH 7.5 + 3 mL 5M NaCl + 12.5 µL 2M spermidine and fulfilled to 50 mL with MilliQwater and one diluted tablet of protease inhibitor)). On a rotator, samples were incubated at room temperature for 1 hour. After the removal of liquid and washing with DIG-300 three times, beads were resuspended in 300 µL Tagmentation buffer (5 mL DIG-300 + 50 µL 1M $MgCl_2$) and incubated at 37°C for 1 hour to perform the tagmentation reaction. After stopping the tagmentation reaction (by adding 10 µL 0.5M EDTA, 3 µL 10% SDS and 2.5 µL 20 mg/mL Proteinase K to each sample), DNAs were released after the lysis of nuclei in the solution at 55°C for one hour., DNA was extracted using a MinElute PCR Purification Kit (Qiagen). Extracted DNA was stored at -20°C. Ultimately, tagged DNA was subjected to amplification with a dual indexing system (Nextera XT Index Kit, Illumina) and purified with SPRI select (Beckman Coulter) twice, as follows: (1) 1.3 volume equivalents of SPRI, and (2) 1.4 volume equivalents of SPRI. DNA libraries' concentration was assessed with Invitrogen Qubit 4 Fluorometer (ThermoFisher Scientific) using Qubit 1X dsDNA HS Assay Kit and a BioAnalyzer 2100 (Agilent Technologies), using Agilent High Sensitivity DNA Kit according to the instructions from manufacturers. Sequencing was performed on an Illumina NextSeq2000 (50 bp, paired-end) at an average depth of 16.17 million reads/library for H3K27ac, H3K27me3, and H3K4me1.

**Cut&Tag data alignment, peak-calling, and analysis.** Cut&Tag data was processed with Cutadapt (4.0) [18] to trim adapter sequences (Nextera Transposase Sequence) with the following parameters '-a CTGTCTCTTATA -A CTGTCTCTTATA -m 25:25' from the 3' end of reads. Duplicated counts were counted by Picard MarkDuplicates (2.23.5) [16]. Against the Mus musculus genome (mm10) reads were mapped using bowtie2 (2.2.6) [19] with default parameters except for '--phred33 --end-to-end --no-mixed --no-discordant -I 10 -X 700 --mm --very-sensitive'. The aligned reads whose mapping quality is less than 10 were filtered by using samtools (1.7) [20]. In addition, reads falling into Encode blacklisted regions v2 [21] were removed using BEDtools intersect (2.30.0) [22]. The Deeptools bamCoverage (3.5.4) [23] was used to generate the BigWig with the following parameters: '-bs 10 -p 10 –normalizeUsing CPM –effectiveGenomeSize 2652783500 –skipNonCoveredRegions –extendReads –ignoreDuplicates'. Peak calling was done on pools of replicates for the histone and the control (naive IgG) libraries with SICER2 (1.0.3) [24] with the parameters 'species mm10, window_size 200, effective_genome_fraction 0.74, false_discovery_rate 0.01, fragment_size= computed on sample with Homer (4.11) [25], gap_size {200, 400, 600, 800, 1000, 2000}, and redundancy_threshold 1'. The peak quality was assessed by the package ChIPQC (1.30.0) [26]. After visual inspection, we selected those with a gap size 200 for H3K4me1 and H3K27ac and 400 for H3K27me3. Detected peaks were combined to get the union of the sex-specific libraries (CPID and Sham for each histone mark) using the tool Bedtools merge (2.30.0) [22]. Then, the number of reads per merged peaks per sample was computed using deeptools multiBamSummary (3.5.5) [23]. Data normalization was performed using the method proposed by Anders and Huber [27] using read counts per peak. PCA was computed on data transformed by a regularized logarithm method calculated with the method proposed by [28]. We further corrected the batch effects for the raw counts per peak sample using sva (3.46.0) [29]. Comparisons of Sham and CPID groups were performed in each sex using DESeq2 (1.38.3) [28] to identify differential peaks (diffPeaks).

**RNA-seq library preparation.** Total RNA was quantified using a NanoDrop Spectrophotometer ND-1000 (NanoDrop Technologies, Inc.) and its integrity was assessed using a 2100 Bioanalyzer (Agilent Technologies). Ribosomal RNAs were depleted from 125 ng of total RNA using QIAseq FastSelect (HMR). cDNA synthesis was achieved with the NEBNext RNA First Strand Synthesis and NEBNext Ultra Directional RNA Second Strand Synthesis Modules (New England BioLabs). The remaining steps of library preparation were done using the NEBNext Ultra II DNA Library Prep Kit for Illumina (New England BioLabs). Adapters and PCR primers were purchased from New England BioLabs. Libraries were quantified using the KAPA Library Quantification Kits - Complete kit (Universal) (Kapa Biosystems). The average size fragment was determined using a LabChip GX II (PerkinElmer) instrument. The libraries were normalized and pooled and then denatured in 0.02N NaOH and neutralized using the HT1 buffer. The pool was loaded at 175pM on a Illumina NovaSeq S4 lane using Xp protocol as per the manufacturer's recommendations. The run was performed for 2x100 cycles (paired-end mode). A phiX library was used as a control and mixed with libraries at 1% level. Program BCL Convert 4.2.4 was then used to demultiplex samples and generate FASTQ reads. The average sequencing depth was 71.47 ± 2.1 million reads/library (mean ± sem).

**RNA-sequencing analysis.** The STAR tool (2.7.10b) [30] was used against the Gencode (GRCm38.p6) reference genome for alignment and quantification (--quantMode). Kept only genes that have at least 10 reads across the 12 samples in each male and female dataset. Read count normalization was done using the Anders and Huber proposed method [28]. The data was analyzed by DESeq2 (1.38.3) with the parameter 'design= ~RIN_scaled+Condition' of DESeqDataSetFromMatrix() function and contrast as CPID vs Sham, where Sham was a reference in the results() function-the differentially expressed genes (DEGs) were identified with a p-value threshold of 0.01.

**Rank-rank hypergeometric overlap analysis.** To visualize the transcriptomic and methylomic profiles in a threshold-free manner, we have used the two approaches: (1) RRHO2 (1.0) [31] for the transcriptomic data and (2) RedRibbon (1.0.1) [32], a package that has been optimized for large datasets, such as, DNA methylation data. For RRHO2, Ensembl gene IDs in each dataset (male and female) were ranked based on their corresponding values of Log2 Fold change and p-values obtained from the function, -log10(P-value) x sign(Log2 Fold Change). Then the RRHO2 function for two datasets was applied with the default parameters (with step size equal to the square root of the list length). For RedRibbon, the whole mm10 genome was partitioned in 500-bp non-overlapping bins. To avoid the sex-bias, we excluded the chromosomes Y and M [33] from binning from male and female data. Whole genome bins were ranked in each sex based on -log10(p-value) x sign(CPID/Sham DNA methylation difference), and were compared using the Redribbon algorithm, which implements an "evolutionary algorithm" for computational efficiency [32]. Hypergeometric p-values were reported for RRHO2 and RedRibbon as -log10(p-values). GO enrichments of genes corresponding to the best hypergeometric p-values were determined by using the enrichGO() function of clusterProfiler package (4.6.2) [34].

**Gene network co-expression Analysis.** We used Multiscale Embedded Gene co-Expression Network Analysis, MEGENA [35], a co-expression approach that identifies a hierarchical organization among co-expressed gene modules [36]. MEGENA was applied to build 3 networks: 1 for transcriptomic data generated in each sex, 1 for the pooled data. Genes from chromosomes X and Y, as well as unknown locations were excluded. Raw counts normalization was performed by the sequencing depth, using the counts per million (cpm) metric, and only genes with at least 1 cpm in at least 80% of the samples were included in network construction. CPM values were then normalized using the Counts adjustment with TMM factor (CTF) approach, as recommended by a recent benchmark [37], followed by correction of RIN effects, using a linear model residualization. Correlations among pairs of genes and Planar Filtered Network (PFN) testing were then computed, followed by clustering of genes into modules (pval <= 0.01; number of genes per module: min, 50; max, half of genes within the dataset). We then determined how each module was enriched for genes identified as DEGs (RNA-seq), or annotated to loci identified during epigenetic differential analyses (CG-DMR, CAC-DMRs, diffLMRs, diffUMRs, diffbins, and diffPeaks for the 3 histone marks), using Fisher's Exact Test (FET) from GeneOverlap (1.38.0) [38]. P-values resulting from these enrichments were then combined using the Stouffer method [39] and the poolr (1.1-1) R package [40]. Preservation of each sex-specific network was assessed against the network built in the other sex using the preservation function of the WGCNA R package [41, 42], which yielded ZSummary values (ZSummary<2, module not preserved; 10>Zsummary>2= moderately preserved; Zsummary>10, strongly preserved).

### 2. Legends of supplementary figures

**Supplementary Figure 1. a.** Evaluation of conversion inefficiency in the CG context using spiked-in Lambda DNA controls. Data are Mean ± SEM. **b.** DNA methylation abundance in the CG and 12 non-CG trinucleotide contexts. Among non-CG contexts, highest levels were observed at CAC, consistent with previous work. A three-way ANOVA showed a significant main effect of trinucleotide context ($F(12, 260)=88,359.43$, $p<2e-16$). **c.** Differentially Methylated Regions (DMR) identified in females in the CG and CAC contexts using 3 algorithms: dmrseq, methylsig, and DSS. **d.** Density plot of ranks of DMR identified in common by DSS and dmrseq (in females, in the CG context). **e.** Manhattan plots of the chromosomic distribution of DMRs in each sex and each cytosine context. **f.** Principal Component Analysis (PCA) of DNA methylation levels at cytosines located within DMRs identified for each sex and cytosine context.

**Supplementary Figure 2. a.** Permutation testing (using regioneR and an expected random distribution) of the significance of observed overlaps among DMRs identified in females and males, in the CG (left panel) or CAC (right) contexts. **b.** Permutation testing (using regioneR and an expected random distribution) of the significance of mean distances observed between nearest neighbours of DMRs identified in females and males, in the CG (left panel) or CAC (right) contexts. **c.** Venn diagrams of overlap among DMRs identified in each sex as a function of chronic pain-induced depression (CPID-effect), or when comparing female and male Sham groups (Sex-effect), in the CG (left) or CAC (right) contexts. JI, Jaccard indexes. **d.** Convergence of chronic pain-induced DNA methylation changes, depicted in males (top) or in females (bottom): overlaps between CG- and CAC-DMRs are shown for genomic positions (left), associated genes (middle), and related Gene Ontology (GO) terms (right).

**Supplementary Figure 3. a.** Threshold-free comparison, using RedRibbon, of CPID-induced DNA methylation changes between males and females, in the CG context. **b.** Threshold-free comparison, using RedRibbon, of CPID-induced DNA methylation changes between males and females in the CAC context. **c.** Venn diagrams of best overlaps identified in each quadrant of RedRibbon analyses from panels a-b, corresponding to genomic bins that showed, in both sexes, a similar increase (upup), or a decrease (downdown), in DNA methylation levels as a function of CPID. JI, Jaccard indexes. **d.** CPID-induced DNA methylation differences observed in each sex: (i) at hyper- or hypo-DMRs that were identified in females or in males, and in the CG or CAC context, and (ii) at Redribbon bins shared between sexes (as of panel c). For example, the left most panel depicts CPID-induced DNA methylation differences observed, in each sex, at the cytosines located within 3 distincts sets of loci: CG-DMRs identified in females; CG-DMRs identified in males; RedRibbon upup bins. **e.** Permutation testing (using regioneR and an expected random distribution) of the significance of mean distances observed between nearest neighbours of RedRibbon bins and DMRs identified in females (right panels) or males (left), in the CG (top panels) or CAC (bottom) contexts.

**Supplementary Figure 4. a.** UpsetR plot of overlaps among the top 10 Transcription Factors (TF) whose binding motifs were found significantly enriched in the 4 categories of DMRs (in the CG or CAC contexts, and in females or males). **b.** Heatmap of the top 10 TFs enriched in the CG- or CAC-DMRs identified in each sex. **c.** Scatter plot of enrichments of Transcription Factor (TF) binding motifs in female CAC-DMRs compared to male CAC-DMRs (Spearman correlation, $r=-0.03$, $p<0.37$).

**Supplementary Figure 5. a.** Representative IGV view of the validation of UMRs and LMRs using H3K4me1 (our data) and H3K4me2 (data from [13]) histone modifications. Males and female data are shown in blue and pink, respectively (MC, Male CPID group; MS, Male Sham; FC, Female CPID; FS, Female Sham). **b.** Profiles of H3K4me1 and H3K4me2 at UMR and LMR regions identified using MethylSeekR. **c.** Violin plots of absolute DNA methylation differences at DMRs, diffLMRs, and diffUMRs. **d.** PCA of DNA methylation levels observed in males at cytosines located within diffLMRs and diffUMRs. **e.** Top 10 GO terms enriched for diffLMRs, diffUMRs, and diffbins.

**Supplementary Figure 6. a.** Convergence of chronic pain-induced DNA methylation changes: overlaps between diffUMRs (top) or diffbins (bottom) identified in females and males are shown for genomic positions (left), associated genes (middle), and related Gene Ontology (GO) terms (right). **b.** PCA of DNA methylation levels observed in males or females at cytosines located within diffbins identified using CluBCpG. **c-e.** UpsetR plots of overlaps between CG-DMRs, CAC-DMRs, diffUMRs, diffLMRs, and diffbins observed for genomic regions (top row), associated genes (middle), and GO terms (bottom), in males (left panels) or in females (right).

**Supplementary Figure 7. a.** Abundance of histone marks according to gene expression, in females. **b.** Gene profiles of histone marks, in males. **c-h.** PCA of the abundance of histone marks at all regions defined as Peaks (see Methods), for each mark and sex.

**Supplementary Figure 8. a.** Piecharts of proportions of diffPeaks identified for each of the 3 histone marks analyzed, in each sex. **b.** Convergence of chronic pain-induced changes in histone marks: overlaps between diffPeaks identified in females and males are shown for genomic positions (left), associated genes (middle), and related Gene Ontology (GO) terms (right). **c.** Heatmaps of histone marks at male CG- and CAC-DMRs (MC, Male CPID; MS, Male Sham).

**Supplementary Figure 9. a.** Heatmaps of histone marks at clusters of CG- and CAC-DMRs identified in males. **b.** Profiles of histone marks for each cluster of male CG-DMR and CAC-DMRs. **c.** Overlaps between females and males of genes associated with cluster-specific CAC-DMRs. **d.** Correlation of transcription factor motif enrichment between female and male cluster 1-DMRs (Spearman correlation, r=0.91, pvalue<1e-300). Similar correlations were observed in the CG context for the 3 clusters of DMR (data not shown). **e.** Boxplot of the distribution of significant TFs enriched in cluster-specific, and the full set, of CAC-DMRs.

**Supplementary Figure 10. a.** Comparison between males and females of enrichment of pain-induced DEGs in GO terms related to the neurovascular system. **b.** Preservation of MEGENA gene coexpression modules identified in one sex when evaluated in the other sex (measured using the preservation function of WGCNA). Each dot represents a module. The colored and labeled dots represent first-generation modules. **c.** Scatter plot of ranks of MEGENA modules determined in each sex by considering multiomic enrichments (using Stouffer p-values, see Main text; r=0.97, p=4.3e-12).

**Supplementary Figure 11. a.** Heatmap of the enrichment of MEGENA modules (shown as rows, ranked using Stouffer p-values in males) for gene sets defining cortical cell-types (columns, using data from [43] generated using the FindMarkers function of the Seurat package). **b.** Heatmap of the enrichment of MEGENA modules (shown as rows, ranked using Stouffer p-values in males) for gene sets defining cortical cell-types (columns, using data from [44]). Abbreviations (retrieved from each publication): ExN/Excit, Excitatory Neurons; InN/Inhib, Inhibitory Neurons; OPC/Oligo_pc, Oligodendrocyte Precursor Cells; ASC/Astro, Astrocytes; MG, Microglia; EC, endothelial Cells; OLG, Oligodendrocytes.

**3. Legends of supplementary tables**

**Supplementary Table 1.** The table contains 4 spreadsheets with the following information: (1) Sequencing depth of Male and Female mice for RNA-seq, EM-seq, and Cut&Tag cohorts. (2) Mean DNA methylation percentage of Male and Female mice for RNA-seq, and EM-seq cohorts. (3) Number of DEGs, DMRs, and diffPeaks identified as a function of CPID. (4). Number of DMRs and diffPeaks identified as a function of sex.

**Supplementary Table 2.** The table depicts the genomic coordinates and descriptive statistics (coverage, DNA methylation levels and differences, direction of change, annotation to gene features) of DMRs identified as a function of CPID in each sex (M, male; F, female) and each cytosine context, or when comparing male and female Sham control groups.

**Supplementary Table 3.** Genes associated with DMRs, diffUMRs, diffLMRs, and diffbins identified as a function of CPID for males (M) or females (F).

**Supplementary Table 4.** GO terms (BPs) associated with DMRs, diffUMRs, diffLMRs, and diffbins as a function of CPID for males (M) and females (F). Reduced terms identified using the rrvgo R package are also shown.

**Supplementary Table 5.** Results from TFmotifView detailing the Transcription Factors enriched for CPID-effect full set of CG-/CAC- DMRs in males and females.

**Supplementary Table 6.** The table depicts the genomic coordinates and descriptive statistics of diffUMRs, diffLMRs, and diffbins identified as a function of CPID in each sex (M, male; F, female).

**Supplementary Table 7.** The table depicts the genomic coordinates and descriptive statistics of diffPeaks identified as a function of CPID in each sex (M, male; F, female) for each histone mark (H3K27ac, H3K27me3, and H3K4me1), or when comparing male and female Sham control groups for each histone mark.

**Supplementary Table 8.** Genes associated with CPID-related diffPeaks for males (M) and females (F).

**Supplementary Table 9.** GO terms (BPs) associated with diffPeaks as a function of CPID for males (M) and females (F). Reduced terms identified using the rrvgo R package are also shown.

**Supplementary Table 10.** The table contains spreadsheets indicating clusters of CPID-related DMRs (see Methods and main text for details).

**Supplementary Table 11.** Results from TFmotifView detailing the Transcription Factors enriched in cluster-specific CG- or CAC-DMRs in males and females.

**Supplementary Table 12.** Results from RNA-sequencing differential expression analysis in males (M) and females (F).

**Supplementary Table 13.** GO terms (BPs) enriched in up- or downregulated DEGs, in males (M) or in females (F).

**Supplementary Table 14.** Lists of genes corresponding to most significant hypergeometric overlaps identified in each quadrant of the RRHO2 analysis when comparing results from RNA-sequencing differential expression analyses between males and females (for example, “uu” corresponds to genes with CPID-induced increased expression in both sexes), along with their enriched GO terms (BP; padj<0.05).

**Supplementary Table 15.** The table contains spreadsheets related to MEGENA results and presenting the following information: (1) CTF normalization values for gene counts used as input for network construction (MS, Male Sham group; MC, Male CPID; FS, Female Sham; FC, Female CPID); (2) Lists of genes belonging to each module, with their number of connections within their module. (3) Enrichment of MEGENA first-generation modules for gene sets associated with CPID-related changes identified for each omic layer, measured using Fisher's Exact Test (FET). Results are shown as p-values for individual omic layer, and as their averaged Stouffer p-values. (4-9) GO analysis of the top 4 modules (m3, m6, m9, m10) and m3 child modules (m32, and m39). Of note, no significant enrichments were identified for m8. (10) Hierarchical relationships among parent/child MEGENA modules, annotated with their respective GO enrichments.

**Supplementary Table 16.** Cell-type enrichments of MEGENA modules based on 3 external single-cell datasets (see main text), assessed using Fisher's exact test.
