## Supplementary Figure 1 for "Sex differences in epigenetic mechanisms of chronic pain-induced depression"

**a**

| Group | Mean ± SEM (Lambda control) |
|---|---|
| Female CPID | 0.004 ± 0.0003 |
| Female Sham | 0.004 ± 0.0004 |
| Male CPID | 0.001 ± 6.7e-05 |
| Male Sham | 0.0008 ± 4.7e-05 |

**b**

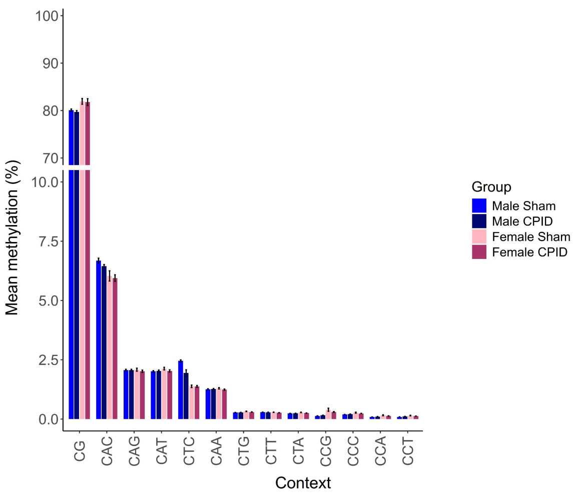


**c**

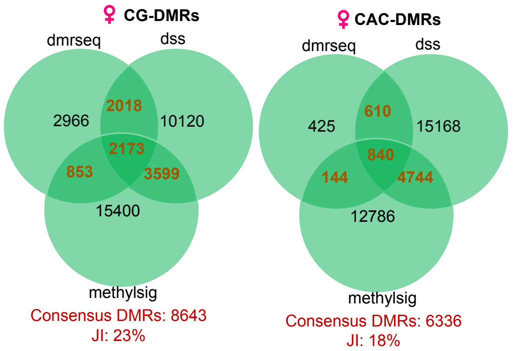


**d**

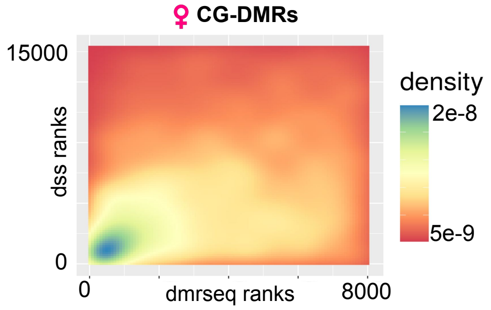


**e**

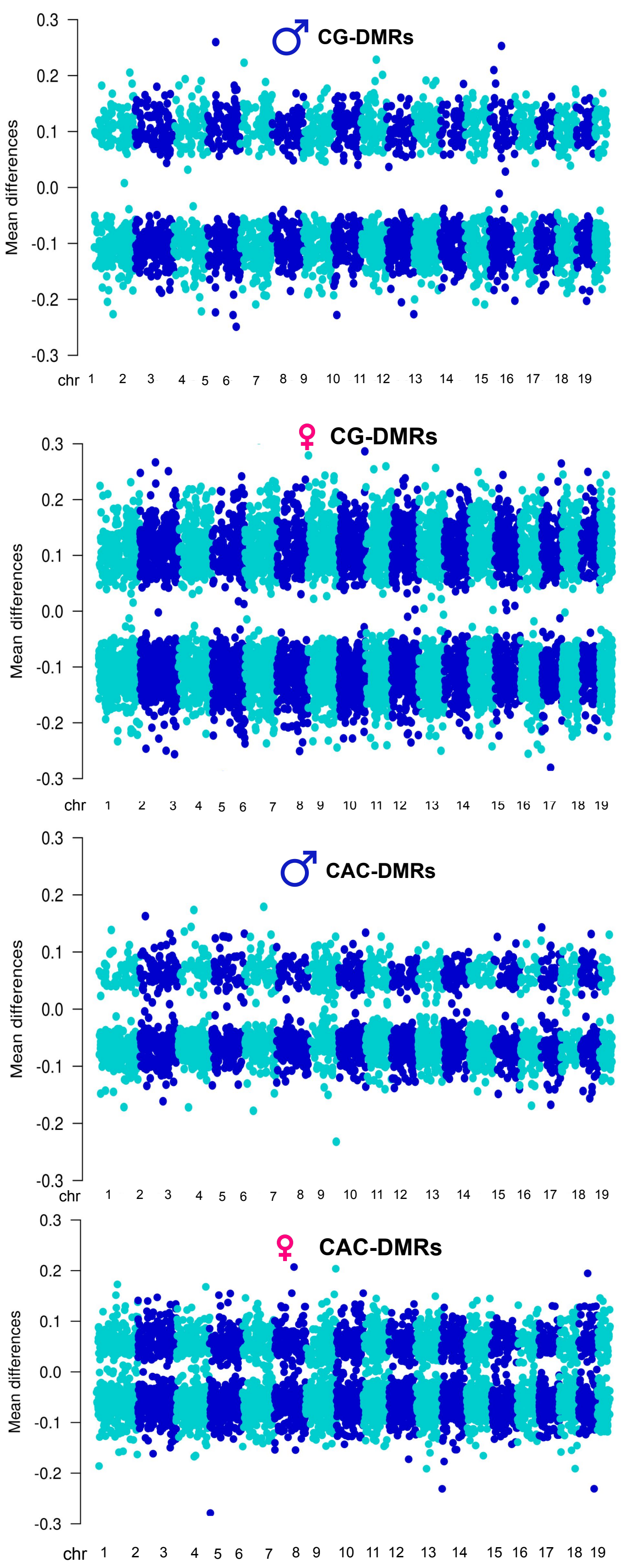


**f**

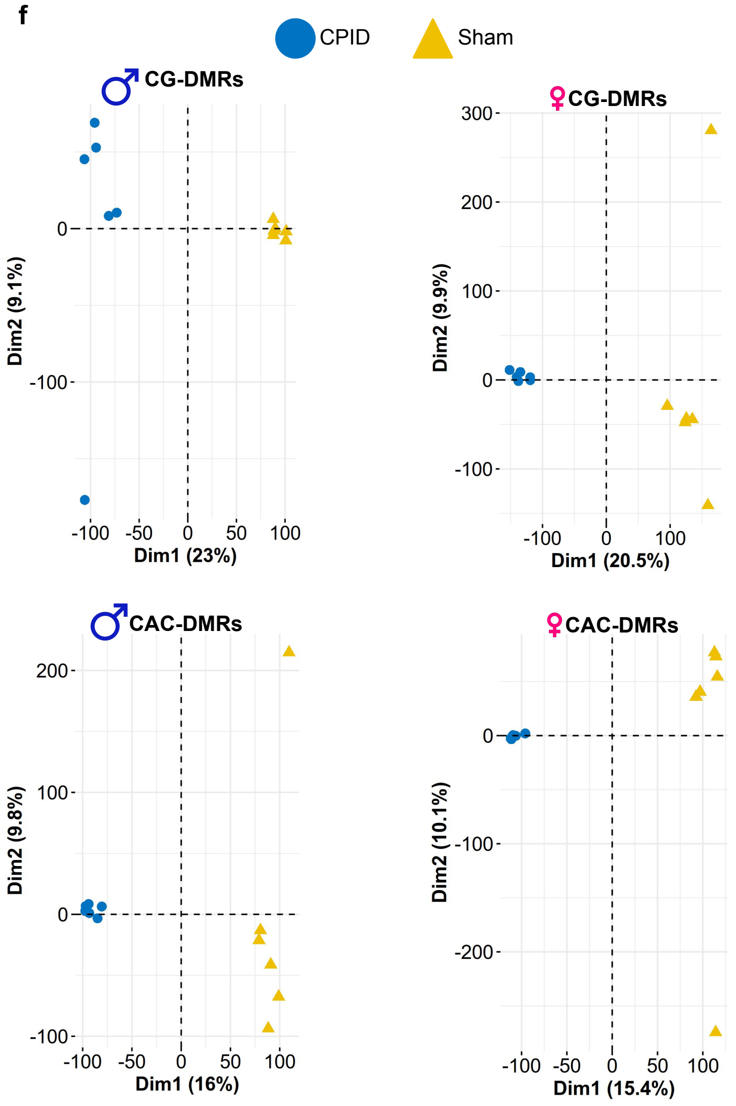
