## Supplementary figures and images for "Sex differences in epigenetic mechanisms of chronic pain-induced depression"

### Supplementary Figure 2

#### Supplementary Figure 2

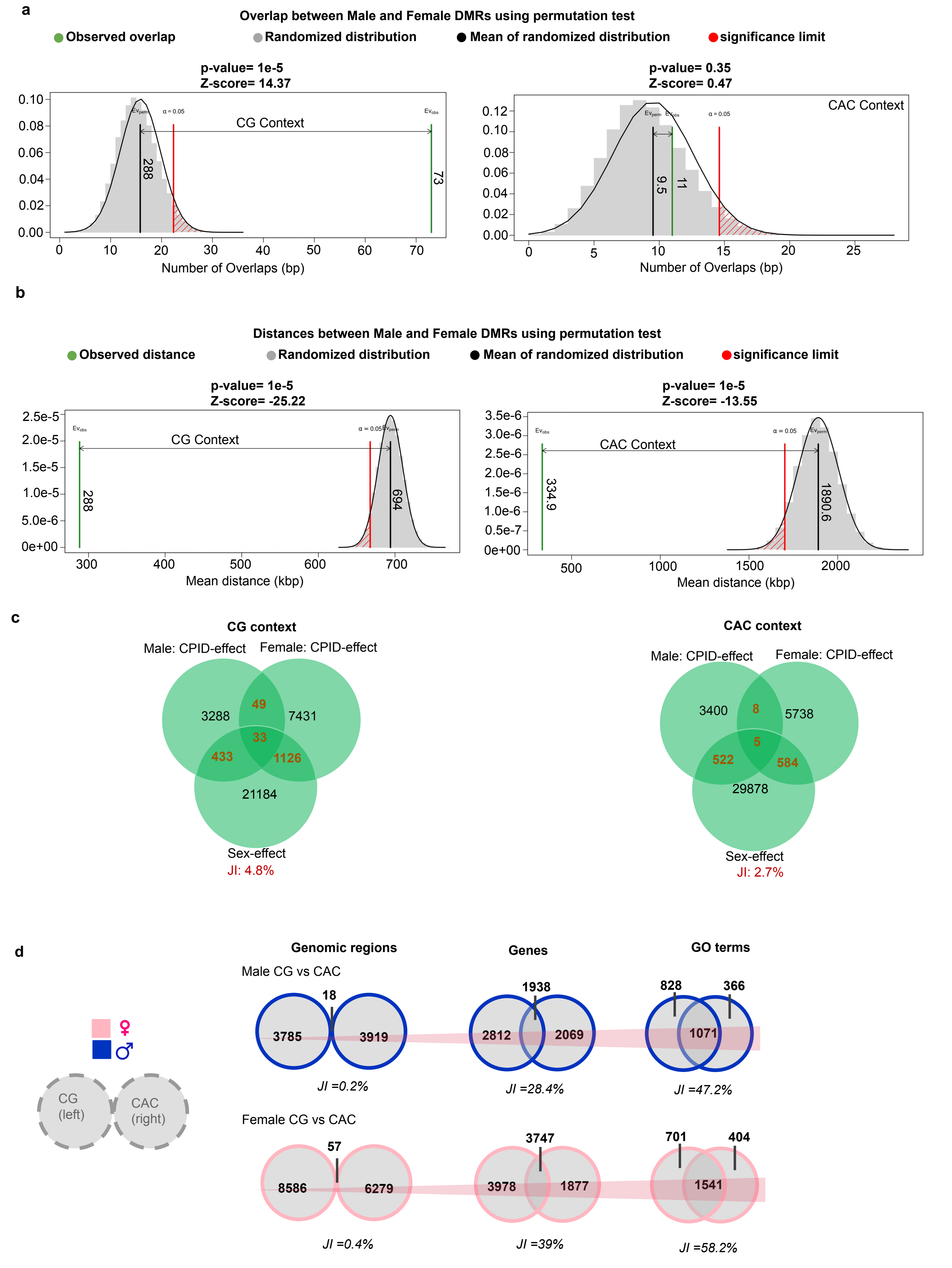

### Supplementary Figure 3

#### Supplementary Figure 3

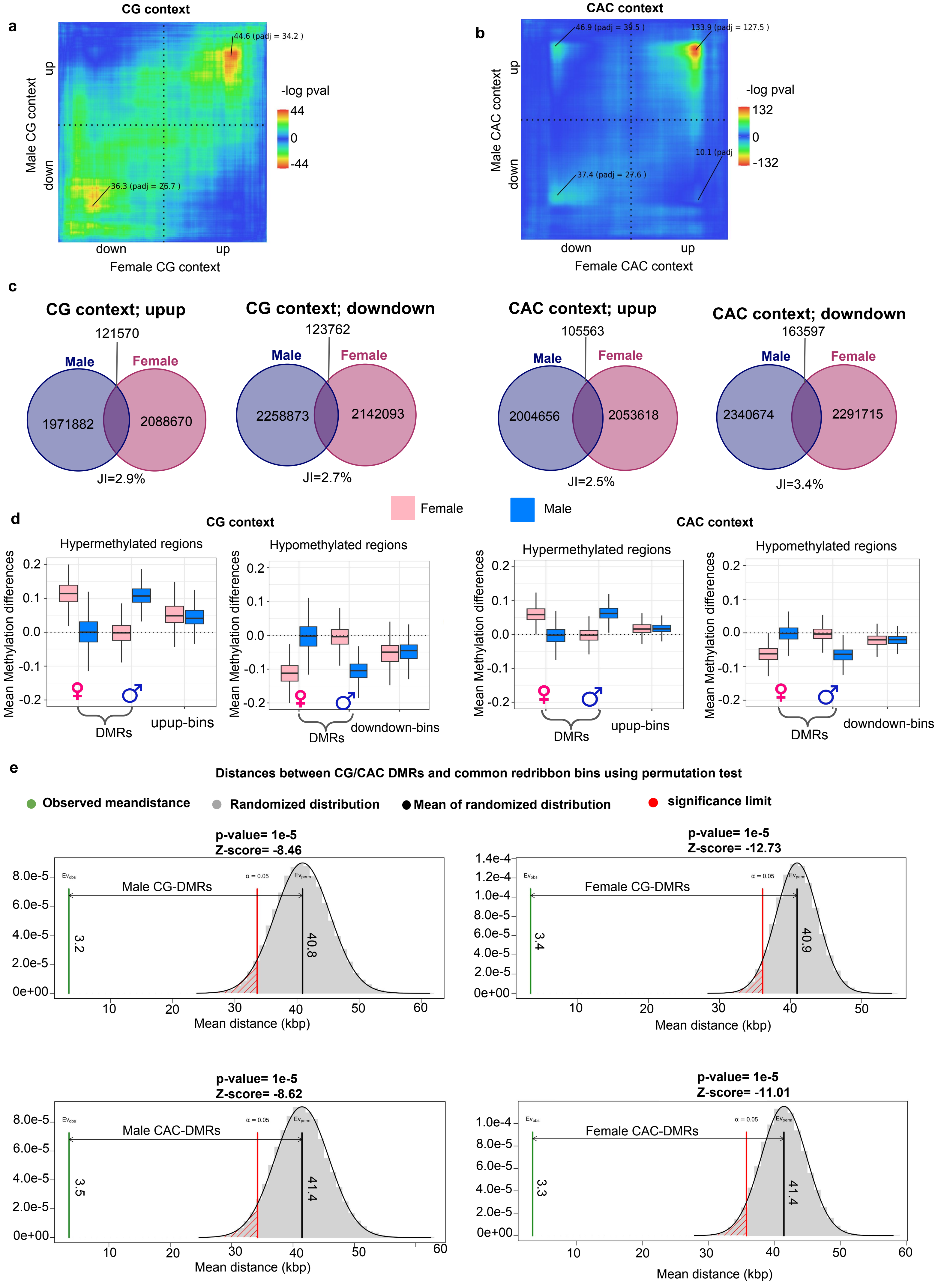

### Supplementary Figure 4

#### Supplementary Figure 4

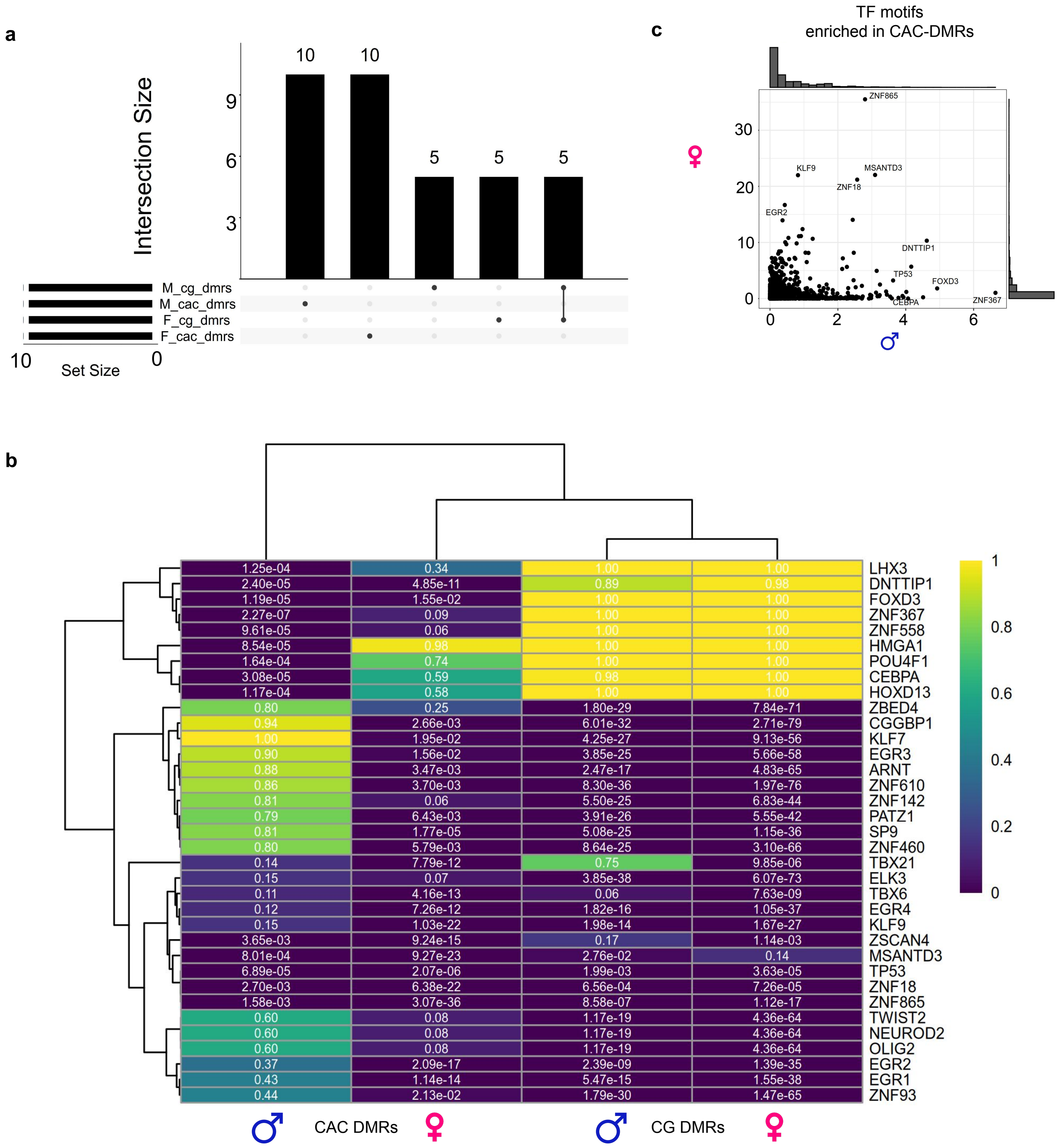

### Supplementary Figure 5

**Supplementary Figure 5**

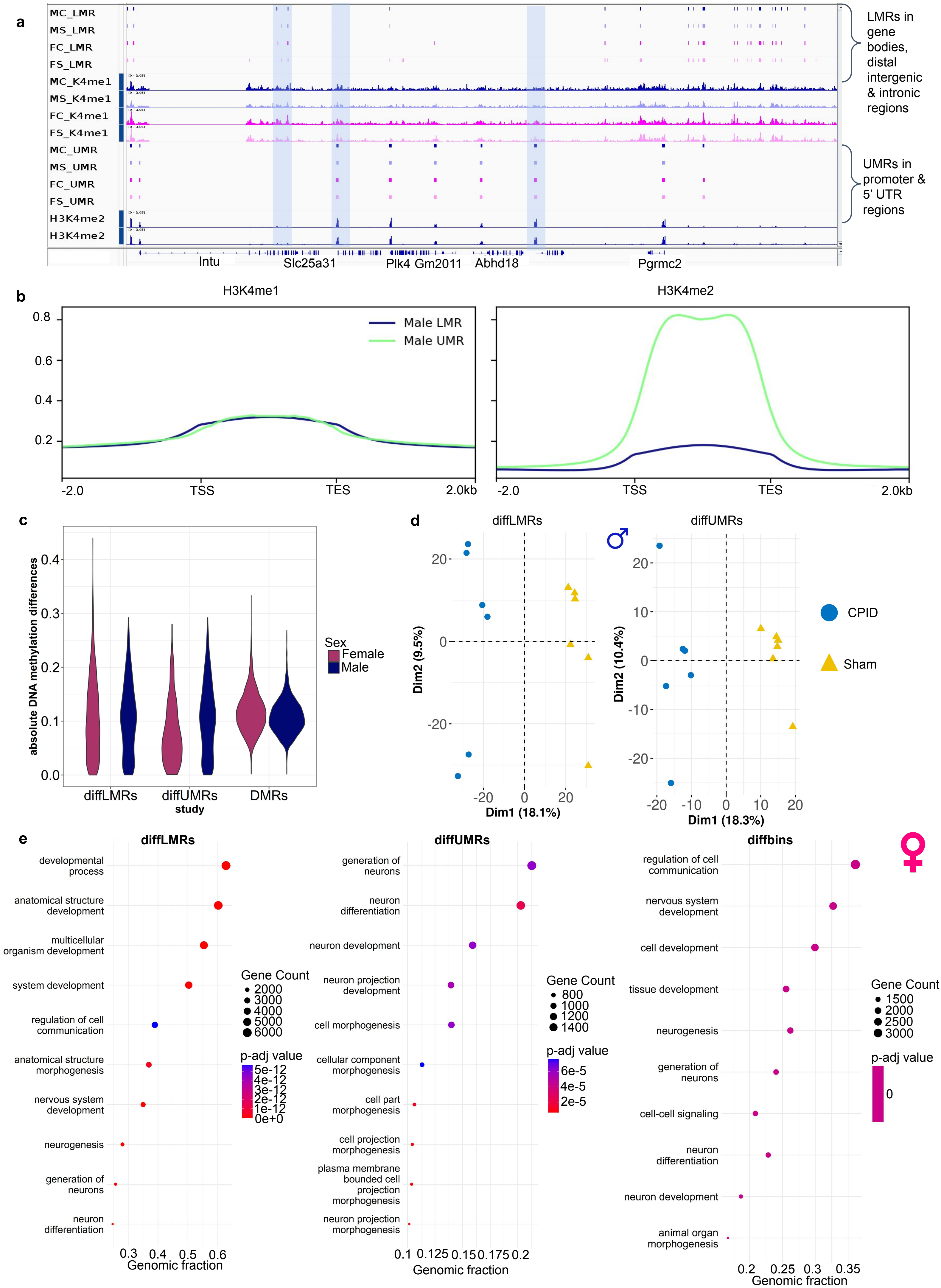

### Supplementary Figure 6

##### Supplementary Figure 6

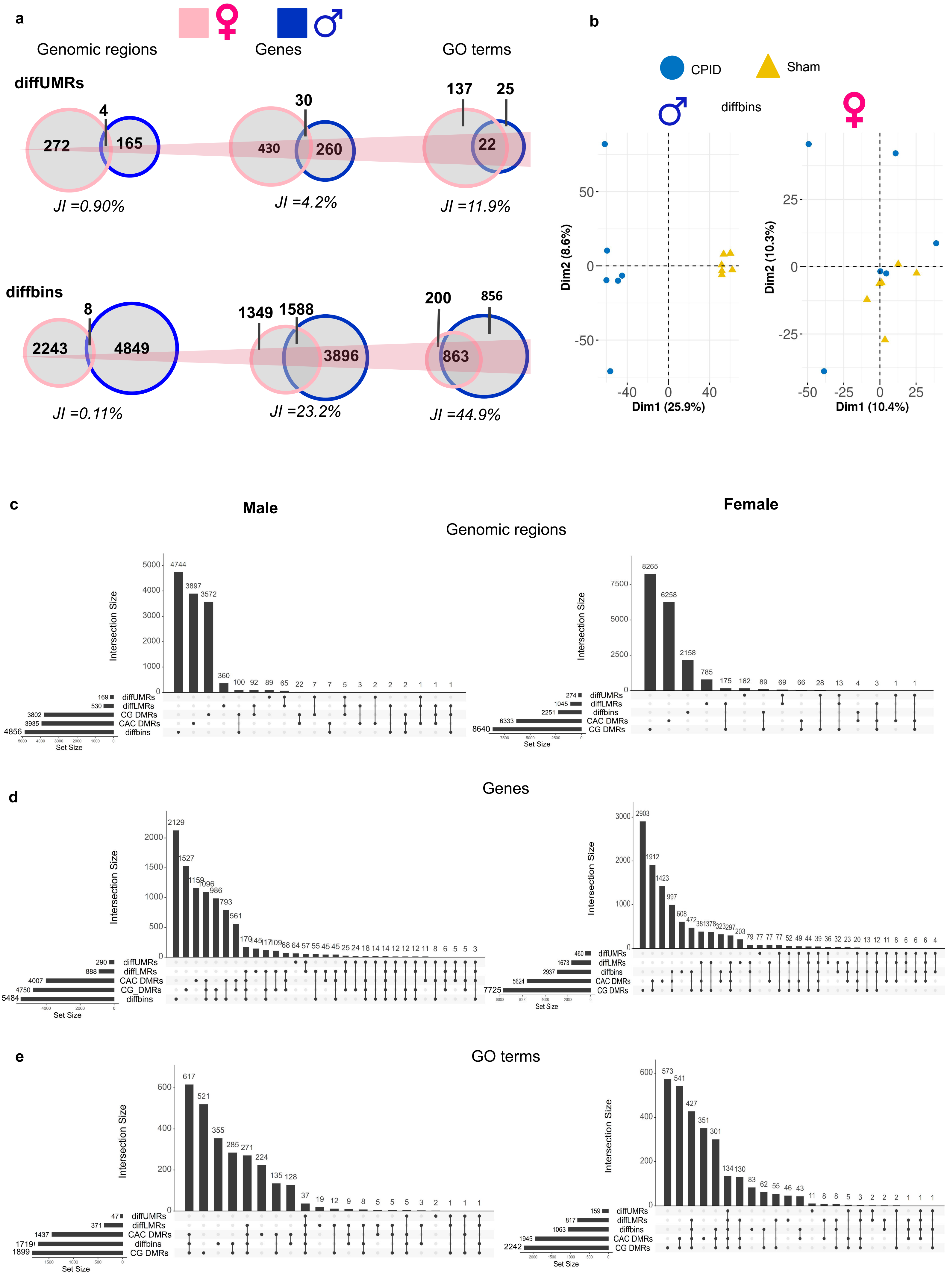

### Supplementary Figure 7

#### Supplementary Figure 7

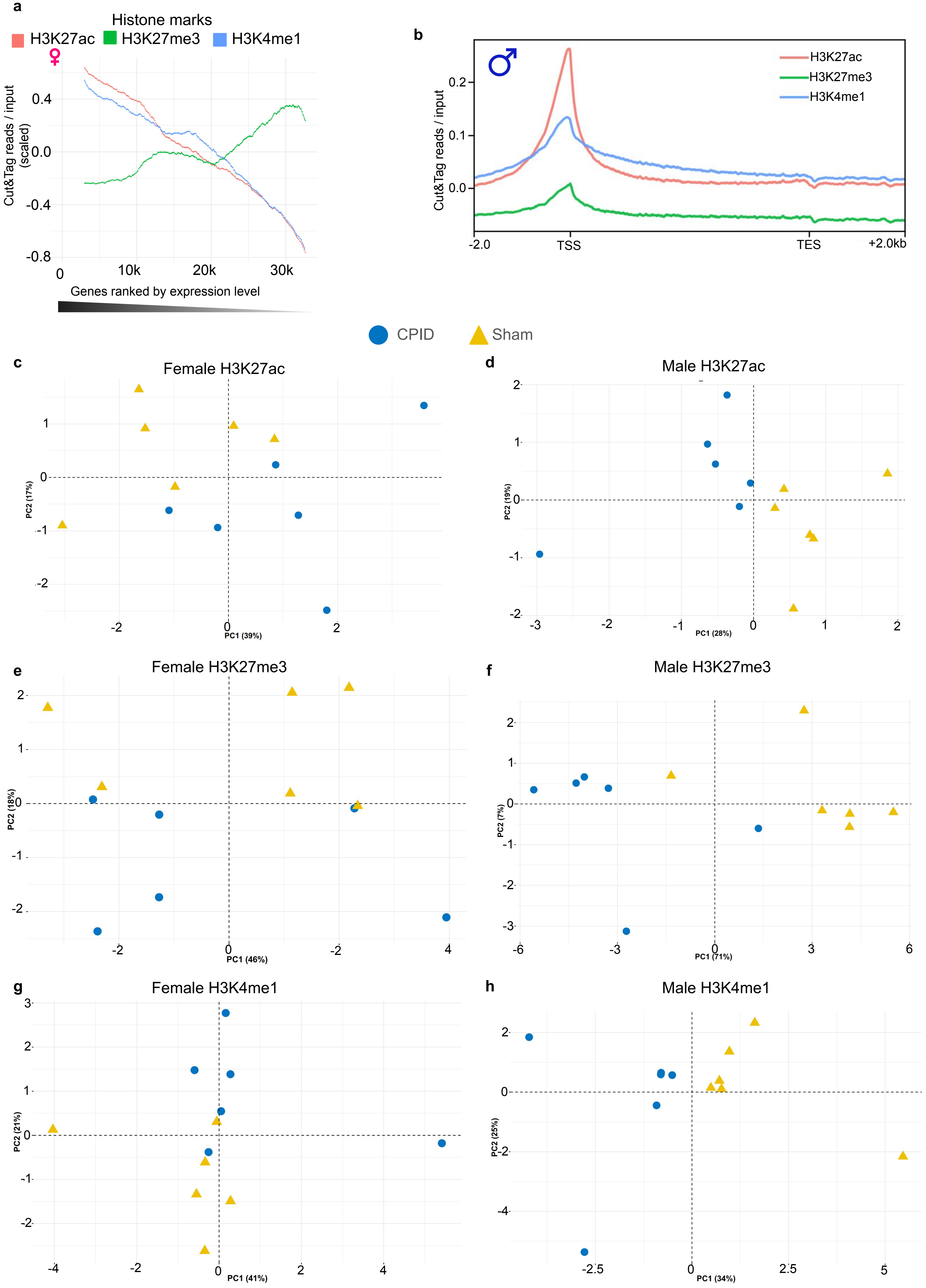

### Supplementary Figure 8

#### Supplementary Figure 8

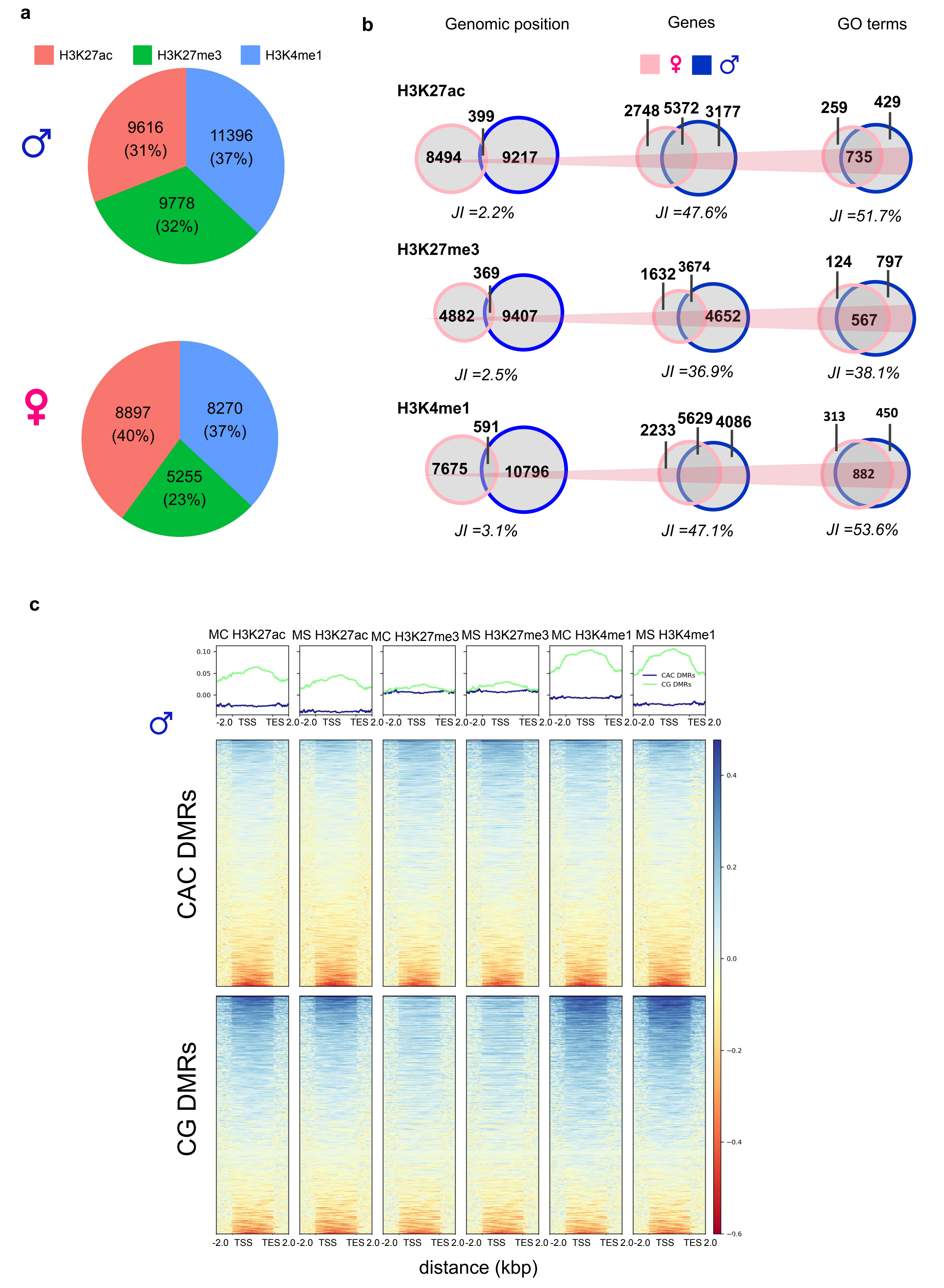

### Supplementary Figure 9

##### Supplementary Figure 9

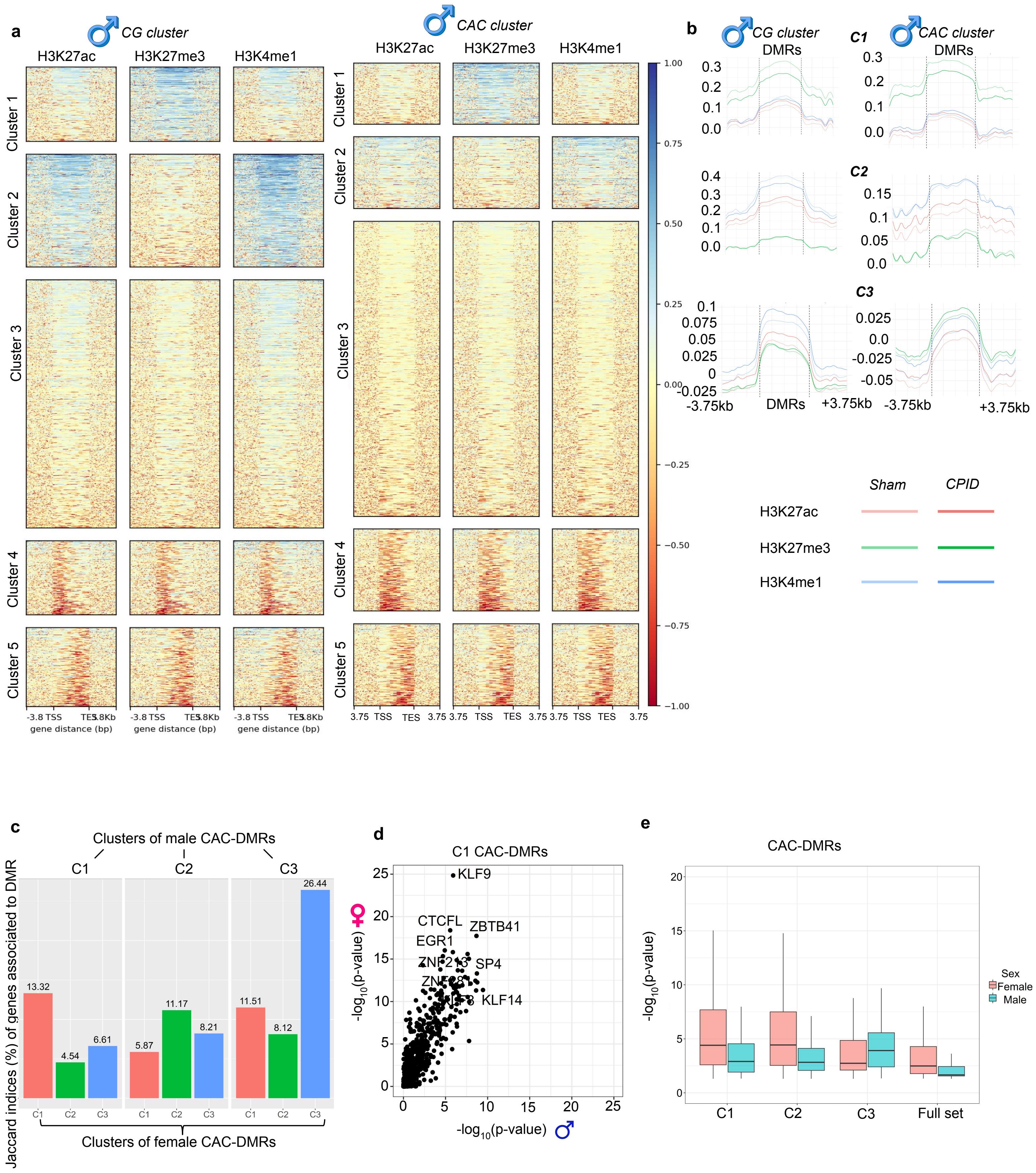

### Supplementary Figure 10

##### Supplementary Figure 10

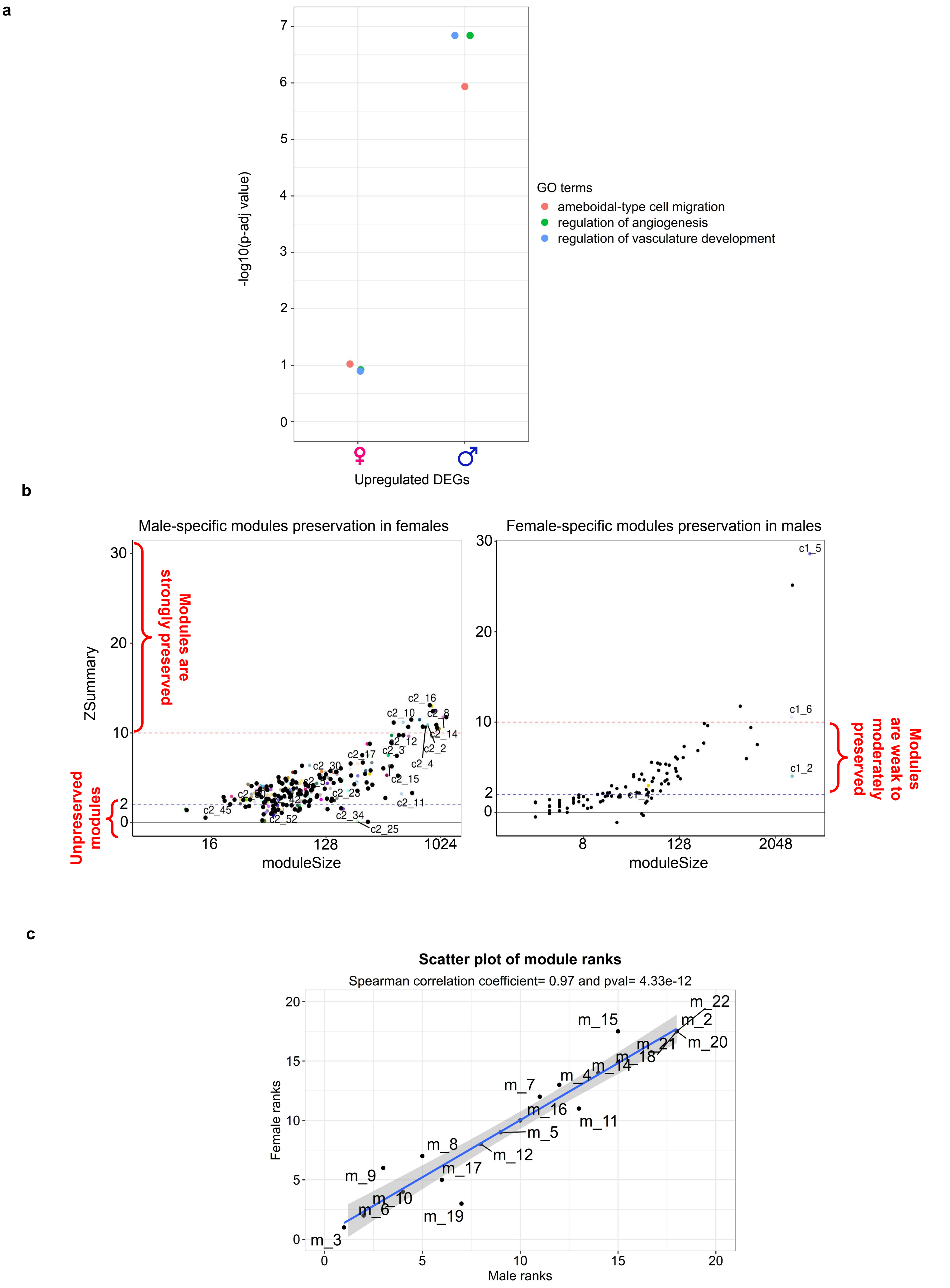

### Supplementary Figure 11

**Supplementary Figure 11**

**a.** Comparison with data from Li et al, 2025

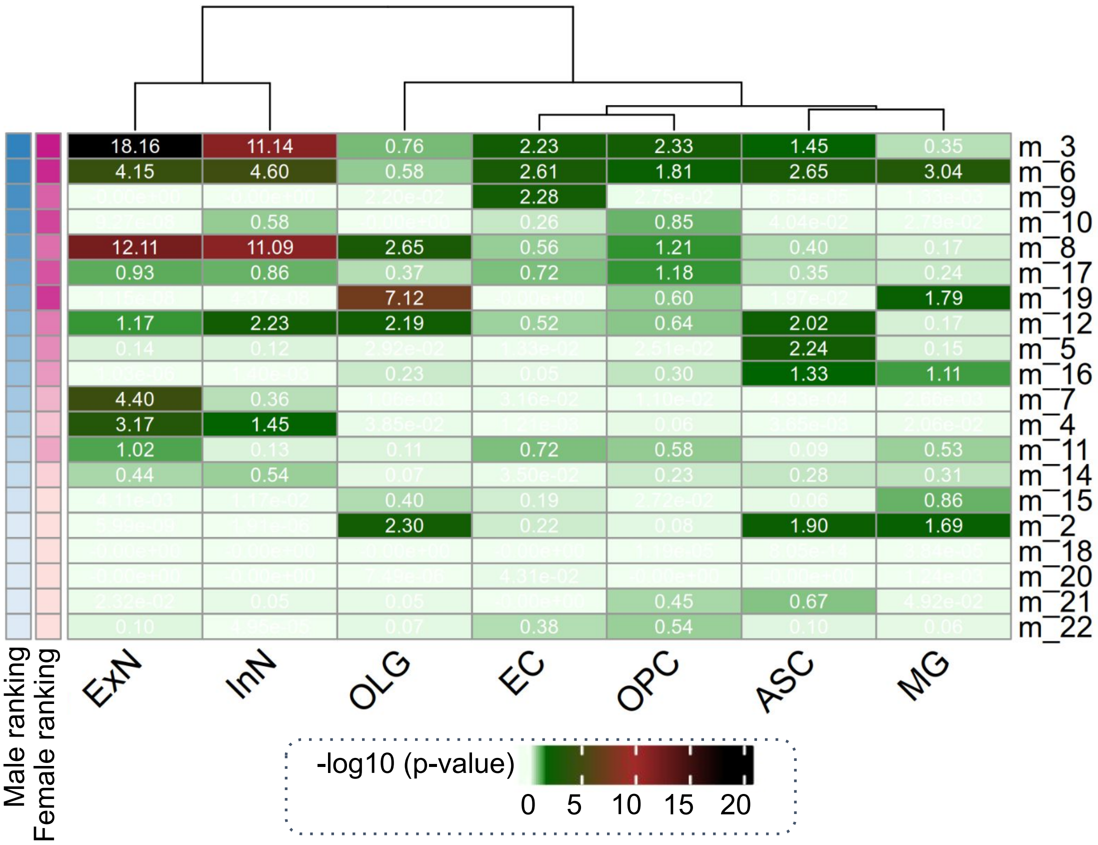


**b.** Comparison with data from Kim et al, 2023

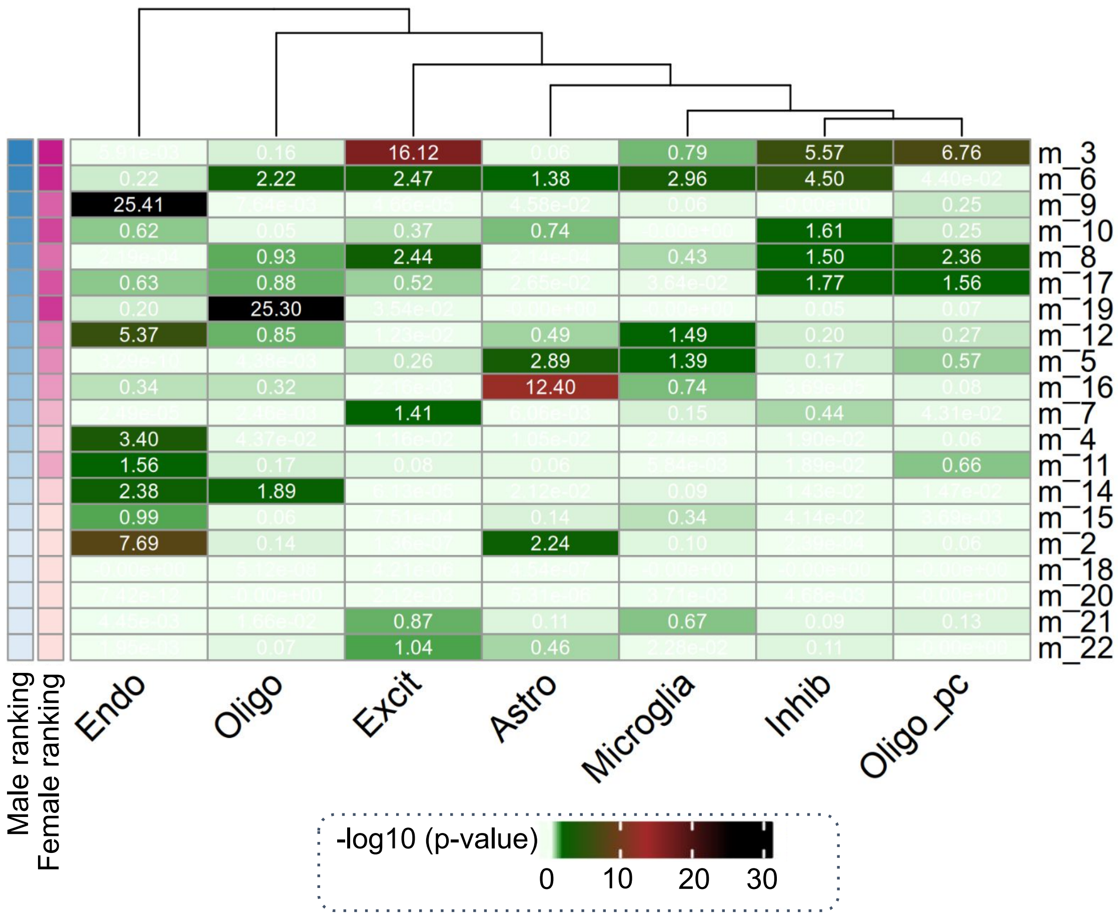
